# Subcellular mRNA localization regulates ribosome biogenesis in migrating cells

**DOI:** 10.1101/829739

**Authors:** Maria Dermit, Martin Dodel, Flora C. Y. Lee, Muhammad S. Azman, Hagen Schwenzer, J. Louise Jones, Sarah P. Blagden, Jernej Ule, Faraz K. Mardakheh

## Abstract

Translation of Ribosomal Protein-coding mRNAs (RP-mRNAs) constitutes a key step in ribosome biogenesis, but the mechanisms which modulate RP-mRNAs translation in coordination with other cellular processes are poorly defined. Here we show that the subcellular localization of RP-mRNAs acts as a key regulator of their translation during cell migration. As cells migrate into their surroundings, RP-mRNAs localize to actin-rich protrusions at the front the cells. This localization is mediated by La-related protein 6 (LARP6), an RNA binding protein that is enriched in protrusions. Protrusions act as hotspots of translation for RP-mRNAs, resulting in enhancement of ribosome biogenesis and overall protein synthesis, which is required for sustained migration. In human breast carcinomas, Epithelial to Mesenchymal Transition (EMT) upregulates LARP6 expression to enhance ribosome biogenesis and support invasive growth. Our findings reveal LARP6 mediated mRNA localization as a key regulator of ribosome biogenesis during cell migration, and demonstrate a role for this process in cancer progression downstream of EMT.

## Introduction

Ribosome biogenesis is elevated in nearly all human cancers^1^. This upregulation is critical for boosting protein synthesis, in order to maintain the increased anabolic demands of malignancy^2^. Enhanced protein synthesis is particularly important for supporting invasion and metastasis in high-grade cancers^3, 4^. Translation of RP-mRNAs has been recognized as the key step in control of ribosome biogenesis ^4, 5^, but how RP-mRNA translation is upregulated in invasive cancers in order to augment ribosome biogenesis remains poorly defined.

It is now clear that rather than being uniformly distributed throughout the cytoplasm, the majority of eukaryotic mRNAs exhibit specific subcellular localizations^6-9^. Such localization can act as a means of localizing the encoded proteins^10^, or function instead as a mechanism for post-transcriptional regulation by modulating the access of mRNAs to different *trans*-acting factors^11-13^. Interestingly, several studies have reported that RP-mRNAs can localize to various forms of cellular projections such as actin-rich protrusions at the front of mesenchymal-like migratory cells or axons of neurons^14-23^. Nevertheless, the molecular mechanism and functional significance of RP-mRNAs localization has remained unclear.

Here we employed a subcellular multi-omics analysis to demonstrate that RP-mRNA localization to actin-rich protrusions is a universal feature of migratory cells. This localization is mediated via LARP6, a microtubule-associated RNA Binding Protein (RBP) that binds to RP-mRNAs to promote their enrichment in protrusions. Protrusions are also highly enriched in translation initiation and elongation factors, acting as hotspots for translation of localized RP-mRNAs. LARP6 dependent localization of RP-mRNAs results in enhancement of ribosome biogenesis and increased global protein synthesis in moving cells, which is required for maintaining their migratory potential. In human breast carcinomas, higher LARP6 protein expression is associated with the invasive mesenchymal-like subtypes. EMT induces LARP6 expression, which acts to promote protein synthesis to support malignant growth and invasion. Crucially, LARP6 regulation of RP-mRNAs localization can be targeted by a small molecule inhibitor which interferes with its RNA binding. Our findings reveal a new mechanism that governs ribosome biogenesis of migratory cells via subcellular localization of RP-mRNAs, and demonstrate a targetable role for this process in cancer downstream of EMT.

## Results

### RP-mRNAs localize to protrusions of all migratory cells

We utilized a micro-porous transwell filter based method^14, 16^ to assess if RP-mRNAs localization to actin-rich protrusions was a conserved feature of all migratory cells. We modified the procedure to allow cells to adhere to the top of the filter first, followed by synchronized induction of protrusion formation through the pores (Fig. 1a). We profiled the subcellular distribution of mRNAs in a diverse panel of normal and malignant migratory human cell-lines from various cell-types and tissues of origin by RNA-sequencing (Fig. 1b). RP-mRNAs were found to be enriched in protrusions of all cell-lines (Fig. 1c & Dataset S1), strongly supporting the notion that their localization to protrusions is universal.

**Fig. 1:**
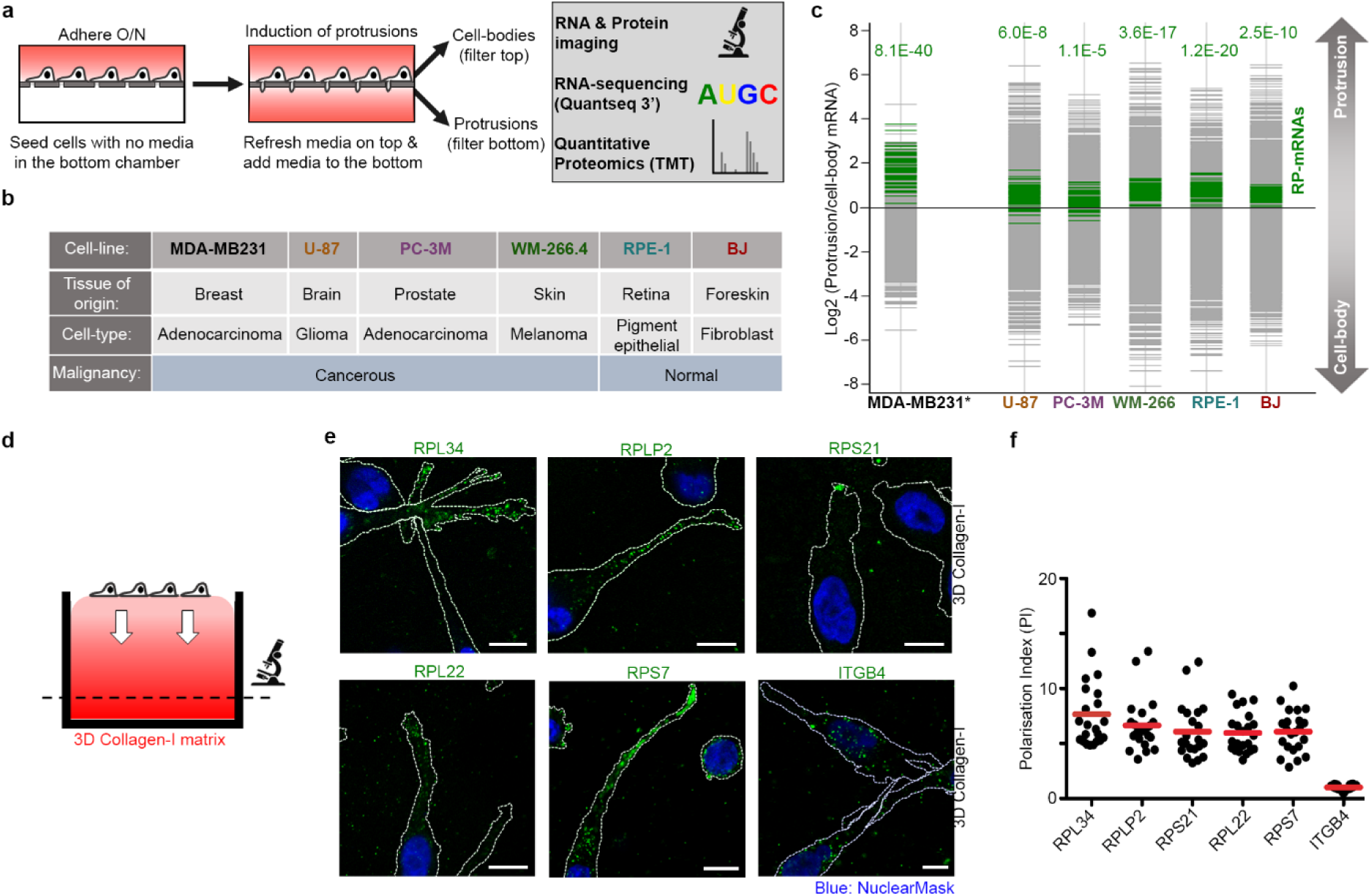
RP-mRNAs universally localize to protrusions. **a**, Schematic representation of transwell based protrusion vs. cell-body analysis experiments. **b**, Panel of normal and malignant cell-lines from diverse tissues of origin, chosen for transwell based profiling. **c**, RP-mRNAs are ubiquitously enriched in protrusions. Transcriptome distributions between protrusion and cell-body fractions in the panel of cell-lines outlined in (b). Benjamini-Hochberg corrected *P-*values of the RP-mRNA enrichments are reported on top of each chart. *MDA-MB231 data was obtained from^16^. **d**, Schematic representation of the experimental setting for RNA-FISH imaging of 3D Collagen-I invading cells. **e**, RP-mRNAs localize to the protrusions of 3D collagen-I invading MDA-MB231 cells. Cell boundaries (dash-lines) were defined by co-staining with anti-tubulin antibody. **f**, Quantification of the polarization index (PI) values^25^ for experiments shown in (e), as a measure of mRNA displacement from the cell-body. Each data point represents a single cell. All scale bars are 10 μm.

Next, we validated our RNA-seq results by RNA-Fluorescence *In Situ* Hybridization (FISH). We used specific RNA-FISH probes against five of the top protrusion enriched RP-mRNAs in the RNA-seq data from MDA-MB231 cells (Dataset S1). All five RP-mRNAs were found to be enriched in protrusions of MDA-MB231 cells while ITGB4 mRNA, which was found to be depleted from protrusions of MDA-MB231 cells by RNA-seq (Dataset S1), did not show an enrichment (Supplementary Fig. 1a-c). To confirm that the observed enrichment of RP-mRNAs is not restricted to transwell settings, we then assessed the localization of RP-mRNAs in actively migrating cells. We chose to assess cell migration in 3D (Fig. 1d), as it is more relevant to the movement of cells *in vivo*^24^. RNA-FISH analysis of MDA-MB231 cells invading through a 3D collagen-I matrix revealed RP-mRNAs to be highly enriched at the tip of cell-protrusions while ITGB4 mRNA remains mostly localized to the perinuclear region (Fig. 1e-f). Collectively, these results suggest that RP-mRNAs localization to protrusions is a conserved feature of mesenchymal-like migrating cells.

### Depletion of LARP proteins reveals a role for LARP6 in RP-mRNAs localization to protrusions

RNA localization is driven by specific RBPs which bind to and mediate transport or anchoring of target transcripts^26^. We therefore hypothesized that specific protrusion localized RBPs must be interacting with and localizing RP-mRNAs to protrusions. As RP-mRNAs localization was conserved across all the cell-lines we tested, localizing RBPs must also be present in protrusions of all of them. To reveal conserved protrusion-localized RBPs, we profiled the distribution of proteins between protrusions and cell-bodies in our panel of cell-lines by tandem mass tagging (TMT) mediated quantitative proteomics^27^ (Fig. 2a & Dataset S2). We then evaluated which known RBPs were enriched in protrusions of all of the cell-lines. 96 RBPs were identified, several of which belong to structurally/functionally related protein categories (Fig. 2a). One such category was the La Related Proteins (LARPs) (Fig. 2a), the prototypical member of which, LARP1, has been previously shown to directly interact with RP-mRNAs through their 5’ Terminal Oligo-Pyrimidine (5’ TOP) motif^28-30^, in a mammalian Target of Rapamycin Complex-1 (mTORC1) dependent manner^31, 32^. However, neither depletion of LARP1 nor inhibition of mTORC1 had an impact on RP-mRNA localization to protrusions (Fig. 2b-c & Supplementary Fig. 2a-c), suggesting that RP-mRNAs localization must be independent of the mTORC1-LARP1-TOP pathway.

**Fig. 2:**
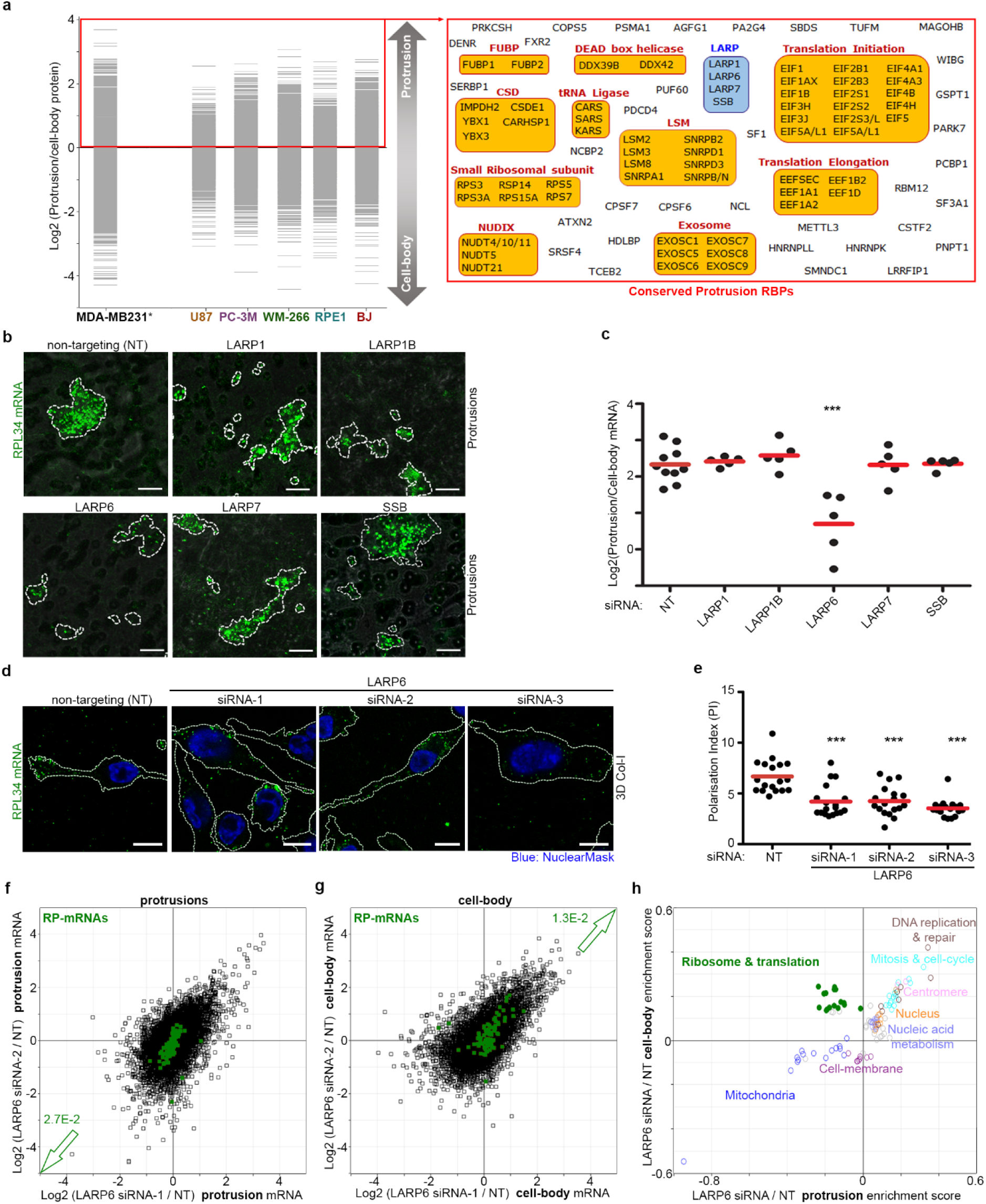
LARP6 localizes RP-mRNAs to protrusions. **a**, Quantitative proteomics reveals conserved protrusion-enriched RBPs. Proteome distributions between protrusions and cell-bodies of the cell-lines listed in Fig. 1B were measured by TMT quantitative proteomics. ‘RNA-binding’ proteins defined according to GOMF database that were present more in protrusions of all cell-lines are listed in the red rectangle. *MDA-MB231 data was obtained from^16^. **b**, siRNA screening reveals LARP6 as a crucial regulator of RP-mRNA localization to protrusions. Representative transwell (grey) RNA-FISH images of RPL34 mRNA (green) in protrusions of MDA-MB231 cells transfected with indicated siRNAs. Cell boundaries (dashed lines) were defined from co-staining with anti-tubulin antibody. **c**, Quantification of RPL34 mRNA enrichment in protrusions from experiments shown in (b). Each data-point represents a large field of view image, pooled from three biological replicates. \*\*\**P*<0.001. **d**, LARP6 depletion prevents RPL34 mRNAs localization to protrusions of MDA-MB231 cells in 3D. Cell boundaries (dash-lines) were defined by co-staining with anti-tubulin antibody. **e**, Quantification of the polarization index values from experiments shown in (d). Each data point represents the PI value for a single cell. \*\*\**P*<0.001. **f**, Depletion of LARP6 reduces RP-mRNA (green) levels in protrusions. Changes in mRNA abundances in MDA-MB231 protrusions were quantified by RNA-seq following LARP6 depletion by 2 independent siRNAs. Benjamini-Hochberg corrected *P*-value of RP-mRNAs decrease is reported next to the arrow. **g**, Depletion of LARP6 increases RP-mRNA (green) levels in the cell-body. Changes in mRNA abundances in MDA-MB231 cell-bodies were quantified by RNA-seq following LARP6 depletion by 2 independent siRNAs. Benjamini-Hochberg corrected *P*-value of RP-mRNAs increase is reported next to the arrow. **h**, RP-mRNAs are mis-localized to cell-bodies upon LARP6 depletions. 2D-annotation enrichment analysis^33^ of data shown in (f) & (g). Each point represents a functional category from GO and KEGG databases, with similar categories being highlighted in the same colors. Upon knockdown of LARP6 with two independent siRNAs, mRNAs coding for ribosomal and translation-related categories (green) change in an anti-correlative fashion, suggestive of mis-localization. All scale bars are 10 μm.

Next, we depleted other LARP family members which were found to be enriched in protrusions of all the cell-lines, along with LARP1B which was found in 5 out of 6 cell-lines (Dataset S2). Only the depletion of LARP6 resulted in a significant decrease in localization of RP-mRNAs (Fig. 2b-c). This decrease was reproduced by 3 independent siRNAs (Supplementary Fig. 2d-f), without having an impact on the ability of the cells to form protrusions per se (Supplementary Fig. 2g), and could be rescued by stable expression of an siRNA resistant GFP tagged LARP6 construct (Supplementary Fig. 2h-i). RP-mRNA localization to protrusions of 3D invading cells was also significantly reduced upon LARP6 depletion (Fig. 2d-e). Lastly, similar to siRNA mediated knockdown of LARP6, short-term (2 hrs) treatment of protruding cells with a small-molecule inhibitor that specifically interferes with LARP6 RNA binding^34^ also reduced RP-mRNA localization to protrusions (Supplementary Fig. 2j-k).

To confirm that the impact of LARP6 depletion was not restricted to just one tested RP-mRNA, we carried out RNA-seq analysis of protrusion and cell-body fractions from control and LARP6 knockdown cells. Depletion of LARP6 by two independent siRNAs resulted in a significant decrease in levels of RP-mRNAs in protrusions along with a concomitant increase in the cell-bodies (Fig. 2f-h; Datasets S3 & S4), strongly suggesting that LARP6 depletion must be resulting in mis-localization of RP-mRNAs from protrusions to cell-bodies.

### Transcriptome-wide iCLIP studies reveal direct binding of LARP6 to RP-mRNAs

We next investigated the localization and function of LARP6. Immunofluorescence (IF) staining revealed endogenous LARP6 to be localized to granules which closely track microtubules (Fig. 3a & Supplementary Fig. 3a-b). In agreement with our proteomics results (Supplementary Fig. 3c), these LARP6 granules were found to be enriched in protrusions (Fig. 3b-c). A fraction of RP-mRNAs co-localized with LARP6 granules, with the co-localization being significantly enhanced in protrusions (Supplementary Fig. 3d-e). Given the co-localization of RP-mRNAs with LARP6 granules in protrusions, we wished to determine whether they directly interact. Collagen type I alpha-1 and alpha-2 (COL1A1 & COL1A2) mRNAs have so far been the only known mRNA partners of LARP6^35-37^. However, COL1A1 and COL1A2 mRNAs were enriched in the cell-bodies of the many cell-lines we examined (Supplementary Fig. 3f), indicating that other mRNA partners are likely to be relevant for the LARP6 function in protrusions. In order to identify direct RNA binding sites of LARP6 across the transcriptome, we utilized MDA-MB231 cells that stably express GFP tagged LARP6 or GFP alone as control (Supplementary Fig. 2h), and performed individual-nucleotide resolution UV crosslinking and immunoprecipitation (iCLIP) (Konig et al., 2010) by anti-GFP beads. In agreement with LARP6 cytoplasmic localization (Fig. 3a), its crosslinking was strongly enriched on exonic compared to intronic regions (Supplementary Fig. 4a-b). We next searched for clusters of LARP6 crosslinking across the genome, which identified 18390 peaks corresponding to likely LARP6 binding sites (Dataset S5). These peaks mapped to a total of 5436 genes (Dataset S6), the vast majority of which were protein coding (Fig. 3d). Category enrichment analysis revealed RP coding transcripts (i.e. ribosome) as the most enriched category (Fig. 3e & Dataset S7), with LARP6 iCLIP peaks found in 73 out of 80 RP-mRNAs (Dataset S6). Other significantly enriched categories included transcripts involved in intracellular trafficking, cell migration, adhesion, and Extracellular Matrix (ECM), among others (Fig. 3e & Dataset S7). Together, these results reveal that LARP6 directly binds a plethora of transcripts, with RP-mRNAs constituting one of its major target mRNA categories.

**Fig. 3:**
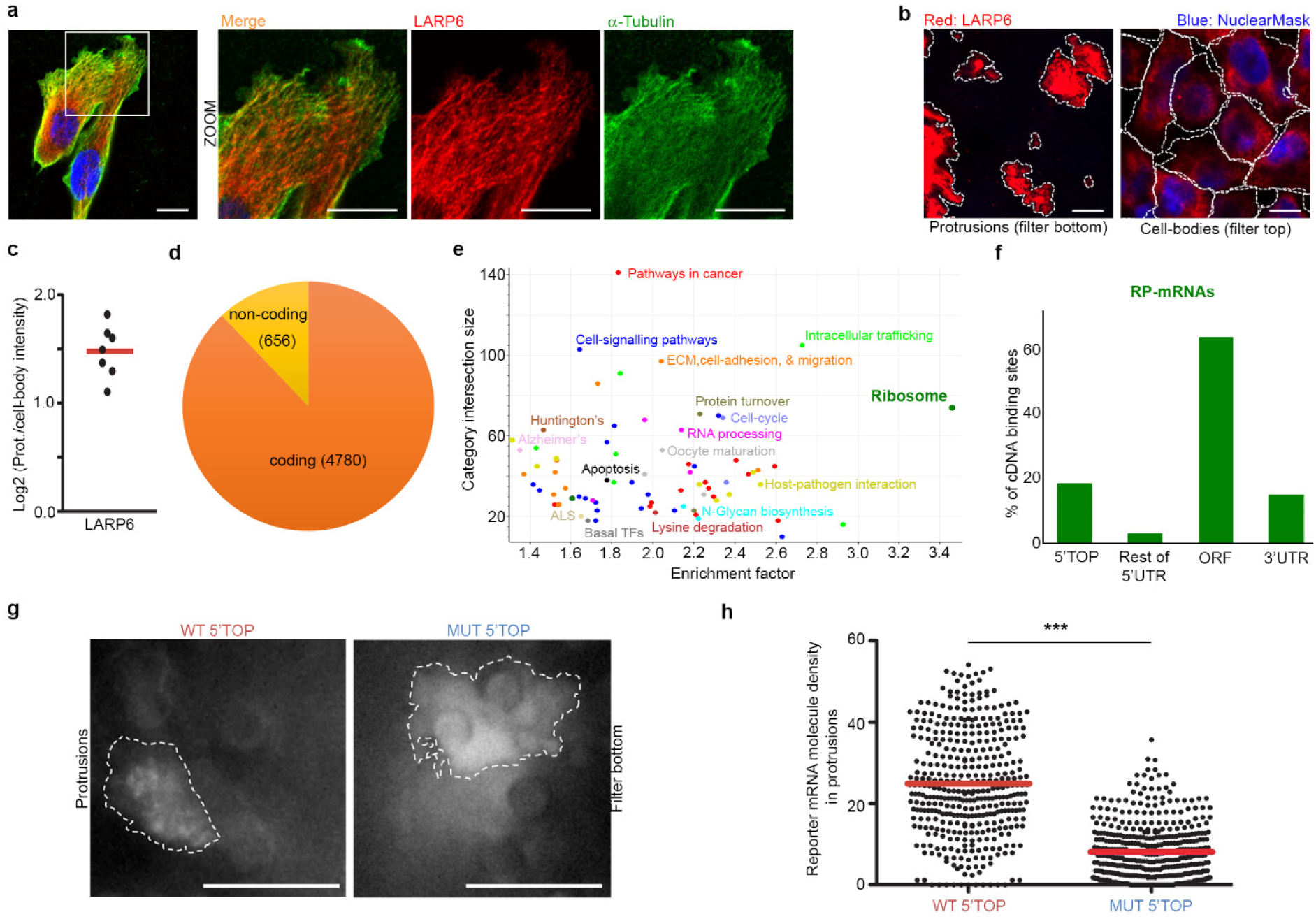
LARP6 is a protrusion enriched RBP which directly binds RP-mRNAs. **a**, LARP6 is localized to granules that track the microtubule cytoskeleton. **b**, LARP6 granules are enriched in protrusions. Representative transwell images of MDA-MB231 cells stained with anti-LARP6 antibody. Cell boundaries (dashed lines) were defined from co-staining with anti-tubulin antibody. **c**, Quantification of LARP6 protrusion to cell-body intensity ratio from experiments shown in (b). Each data-point represents a large field of view image. **d**, Prevalence of coding vs. non-coding RNAs amongst LARP6 direct binding targets. **e**, Fisher’s exact test analysis (FDR < 0.02) of mRNA categories that are significantly over-represented amongst the identified LARP6 targets, with similar categories being highlighted in similar colors. The KEGG category of Ribosome (green), comprised of RP-mRNAs, is significantly enriched amongst LARP6 binding targets. **f**, LARP6 interacts with RP-mRNAs via multiple regions. Distribution of LARP6 binding regions amongst RP-mRNAs. **g**, WT 5’TOP motif is sufficient for RP-mRNA localization to protrusions. Representative still images of the GFP-MCP signal in transwell protrusions of WT or MUT 5’TOP reporter^38^ expressing MDA-MB231 cells as described in Supplementary Fig. 4d, showing a punctate pattern indicative of association with single mRNA molecules in protrusions of WT but not MUT 5’TOP reporter expressing cells. **h**, Quantification of distinct single molecule reporter mRNA counts in experiments shown in (g). Single frames from a total of 25 (WT) and 28 (MUT) time-lapse videos, each taken for 3 seconds at 0.2 second intervals, were quantified. \*\*\**P*<0.001. All scale bars are 10 μm.

We then investigated the mechanism of LARP6 binding and regulation of RP-mRNAs. Around 60% of iCLIP peaks within RP-mRNAs were located in the ORF, with the remaining peaks mainly mapping to the 5’TOP motif, followed by the 3’UTR, and a minor proportion to regions downstream of 5’TOP in the 5’UTR (Fig. 3f). We also detected LARP6 peaks within introns of 43 RP genes, but the majority of these peaks fully overlapped with 37 of the annotated Small nucleolar RNAs (SNORs) that are encoded within the introns of RP genes (Supplementary Fig. 4c & Dataset S6). The positioning of these peaks indicates that LARP6 also binds to SNORs that are processed from the introns of RP-mRNAs. Most RP-mRNAs contained multiple LARP6 iCLIP peaks mapping to different regions (Supplementary Fig. 4c). Nevertheless, using an MS2 based live-cell RNA imaging system^39^ (Supplementary Fig. 4d-e & Videos S1-4), we could show that a single 5’TOP motif results in mRNA localization to protrusions (Fig. 3g-h & Videos S5-6). Together, these results reveal that LARP6 binds to RP-mRNAs via multiple modes, with 5’TOP binding being sufficient to target mRNAs to protrusions.

### LARP6-dependent RP-mRNA localization enhances RP synthesis and ribosome biogenesis

Next, we investigated the functional consequence of RP-mRNA targeting to protrusions. Our profiling of protein distributions between protrusions and cell-bodies had revealed many translation initiation and elongation factors as ubiquitously enriched in protrusions (Fig. 2a). In fact, time-course analysis of the proteome distribution between protrusions and cell-bodies of MDA-MB231 cells showed that proteins involved in translational initiation and elongation accumulate in protrusions early on (Supplementary Fig. 5a-b, and Dataset S8). We therefore hypothesized that this enrichment could lead to higher local levels of translation, making protrusions function as hotspots for protein synthesis in migrating cells. To assess this hypothesis, we mapped the subcellular distribution of translation sites in MDA-MB231 cells using RiboPuromycylation^40^. We optimized the RiboPuromycylation method so that it could be used concurrently with RNA-FISH (Supplementary Fig. 5c). In agreement with the observed accumulation of translation initiation and elongation factors, time-course RiboPuromycylation analysis revealed translation sites to be enriched in protrusions (Supplementary Fig. 5d). Moreover, co-localization of RP-mRNAs with translation sites was significantly higher in protrusions than the cell-bodies (Supplementary Fig. 5e), suggesting that localized RP-mRNAs are likely to undergo more translation. Indeed, using a pulsed-SILAC^41^ based strategy (Fig. 4a), we could show that translation of RPs was significantly enhanced after protrusion formation (Fig. 4b-c, & Dataset S9-10). Importantly, both basal as well as protrusion induced nascent RPs strongly accumulated in the nucleus following their translation (Supplementary Fig. 6a-c & Dataset S11), suggesting that similar to the basally translated RPs, most protrusion synthesized RPs travel back to the nucleus to participate in canonical ribosome biogenesis^42^.

**Fig. 4:**
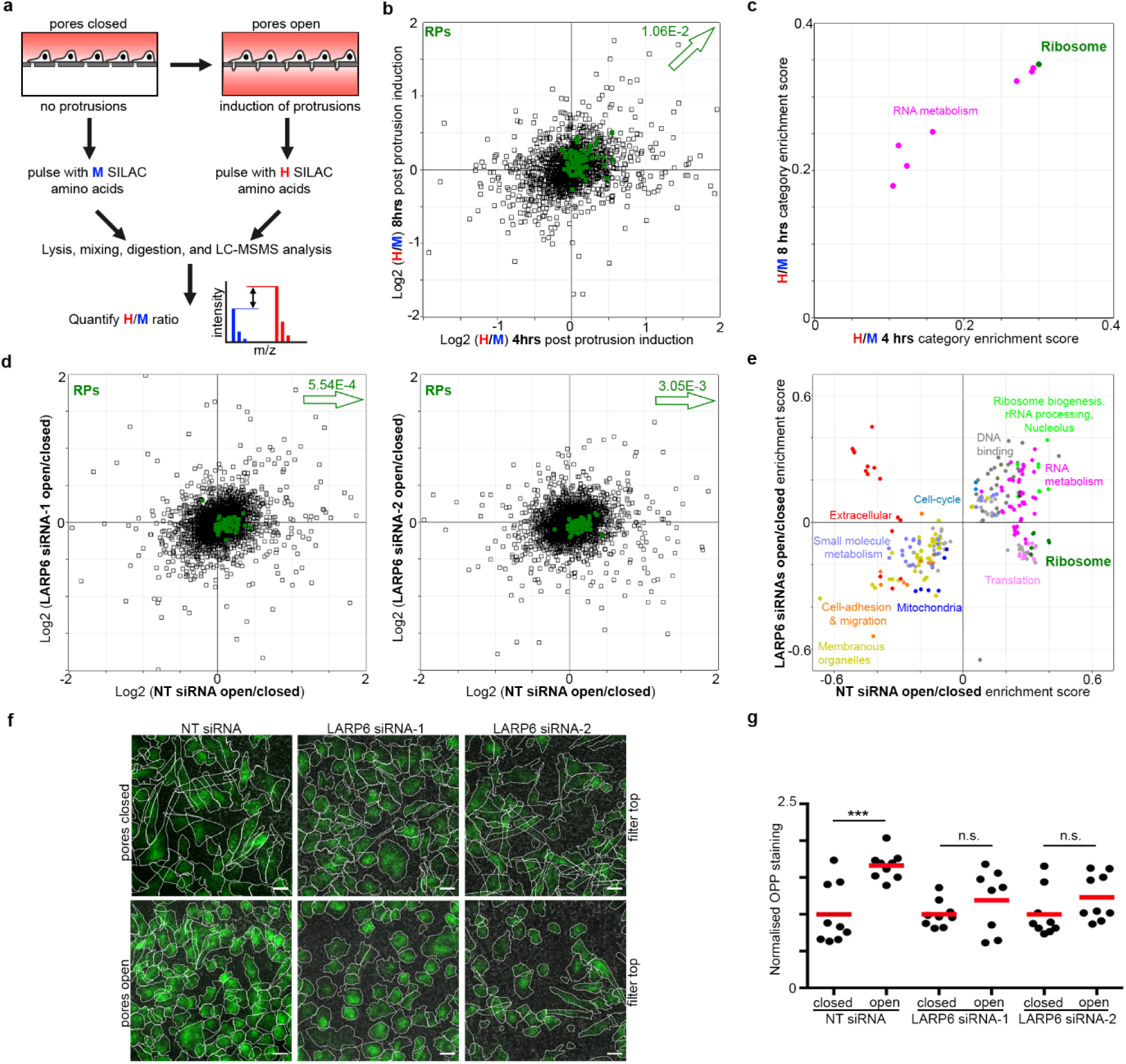
Protrusion localization of RP-mRNAs enhances their translation and ribosome biogenesis. **a**, Schematic diagram of the pulsed-SILAC analysis of translation rate changes induced by protrusion formation. **b**, Protrusion formation in MDA-MB231 cells enhances the translation of RPs (green), as measured according to (a). Benjamini-Hochberg corrected *P*-value for RPs increase is reported next to the arrow. **c**,. 2D-annotation enrichment analysis of data shown in (b). Each data-point represents a functional category from GO and KEGG databases, with similar categories being highlighted in the same colors. The KEGG Ribosome category, comprised of all RPs, is the highest translationally induced category. **d**, Protrusion formation enhances RP (green) levels in a LARP6 dependent manner. Changes in the proteome of MDA-MB231 cells upon protrusion induction for 24 hrs were quantified by TMT mediated proteomics in NT control vs. two independent LARP6 siRNA depleted cell populations (left and right panels). Benjamini-Hochberg corrected *P*-values of the shifts in RP levels are reported next to the arrows. **e**, 2D-annotation enrichment analysis of data shown in (d). Each data point represents a protein category from GO and KEGG databases, with similar categories highlighted by similar colors. Ribosome protein categories comprised of all RPs (dark green) show upregulation upon protrusion induction in the control but not LARP6 depleted cells. **f**, Protrusion formation enhances overall protein synthesis in a LARP6 dependent manner. MDA-MB231 cells treated with indicated siRNAs were seeded on transwells and either prevented from protruding through pores (pores closed), or allowed to form protrusions (pores open) for 24 hrs, before OPP labelling (green). Representative images of the cells from top of the filters are displayed. Scale bars are 20 μm. **g**, Quantification of OPP levels from experiments shown in (f). n.s.: non-significant; \*\*\**P*<0.001.

Whilst augmented translation of RP-mRNAs is necessary for increased ribosome biogenesis, newly synthesized RPs are normally degraded if not incorporated into new ribosomes^43^. We therefore wanted to test whether enhanced translation of RP-mRNAs upon protrusion induction does indeed result in higher stable levels of RPs. TMT mediated quantitative proteomics revealed that whilst short-term (2 hours) protrusion formation did not significantly change total RP levels, long-term (24 hours) formation resulted in a significant increase in total RP levels, along with other protein categories involved in ribosome biogenesis (Supplementary Fig. 6d-e & Dataset S12-13). Accordingly, O-propargyl-puromycin (OPP) labelling^44^ revealed a significant boost in the overall protein synthesis following long-term protrusion formation (Supplementary Fig. 6f-g). Importantly, depletion of LARP6 specifically inhibited upregulation of RP levels following long-term protrusion formation (Fig. 4d-e & Dataset S14-15). Enhancement of overall protein synthesis upon protrusion formation was also inhibited by LARP6 depletion (Fig. 4f-g). Together, these results demonstrate that formation of protrusions and the ensuing LARP6-dependent localization of RP-mRNAs promotes RP synthesis and ribosome biogenesis, leading to enhanced overall protein synthesis in migrating cells.

### LARP6 is important for ribosome biogenesis in highly migratory cells

We next investigated whether LARP6 contributes toward a significant proportion of RP synthesis in highly migratory MDA-MB231 cells. Using SILAC, we quantified the impact of LARP6 depletion on the proteome of actively growing MDA-MB231 cells (Fig. 5a). Levels of RPs were significantly decreased upon LARP6 knockdown (Fig. 6b & Dataset S16). Category enrichment analysis revealed that RPs were amongst the most downregulated protein categories (Supplementary Fig. 7a & Dataset S17). As availability of RPs is crucial for the processing and maturation of rRNAs during ribosome biogenesis, a substantial decrease in their expression should result in accumulation of otherwise transient pre-rRNA transcripts^45^. Accordingly, depletion of LARP6 resulted in a significant accumulation of pre-rRNAs that contain the 5′ External Transcribed Spacer (5’ETS) (Fig. 5c), suggesting that the decrease in RP levels upon LARP6 depletion must be significant enough to hamper rRNA processing.

We next assessed whether depletion of LARP6 compromised migration of invasive cancer cells. LARP6 knockdown significantly reduced the ability of MDA-MB231 cells to invade through 3D Collagen (Fig. 5d-e). LARP6 knockdown also decreased the viability of MDA-MB231 cells, but this decrease was only significant after longer-term depletion of LARP6 (Supplementary Fig. 7b), suggesting that the observed decrease in migration is not due to loss of viability. Accordingly, LARP6 knockdown significantly affected the long-term growth of MDA-MB231 cells as revealed by clonogenic assays (Fig. 5f-g). Together, these results suggest that in highly motile cells, migration and proliferation is reliant on on LARP6 dependent ribosome biogenesis.

**Fig. 5:**
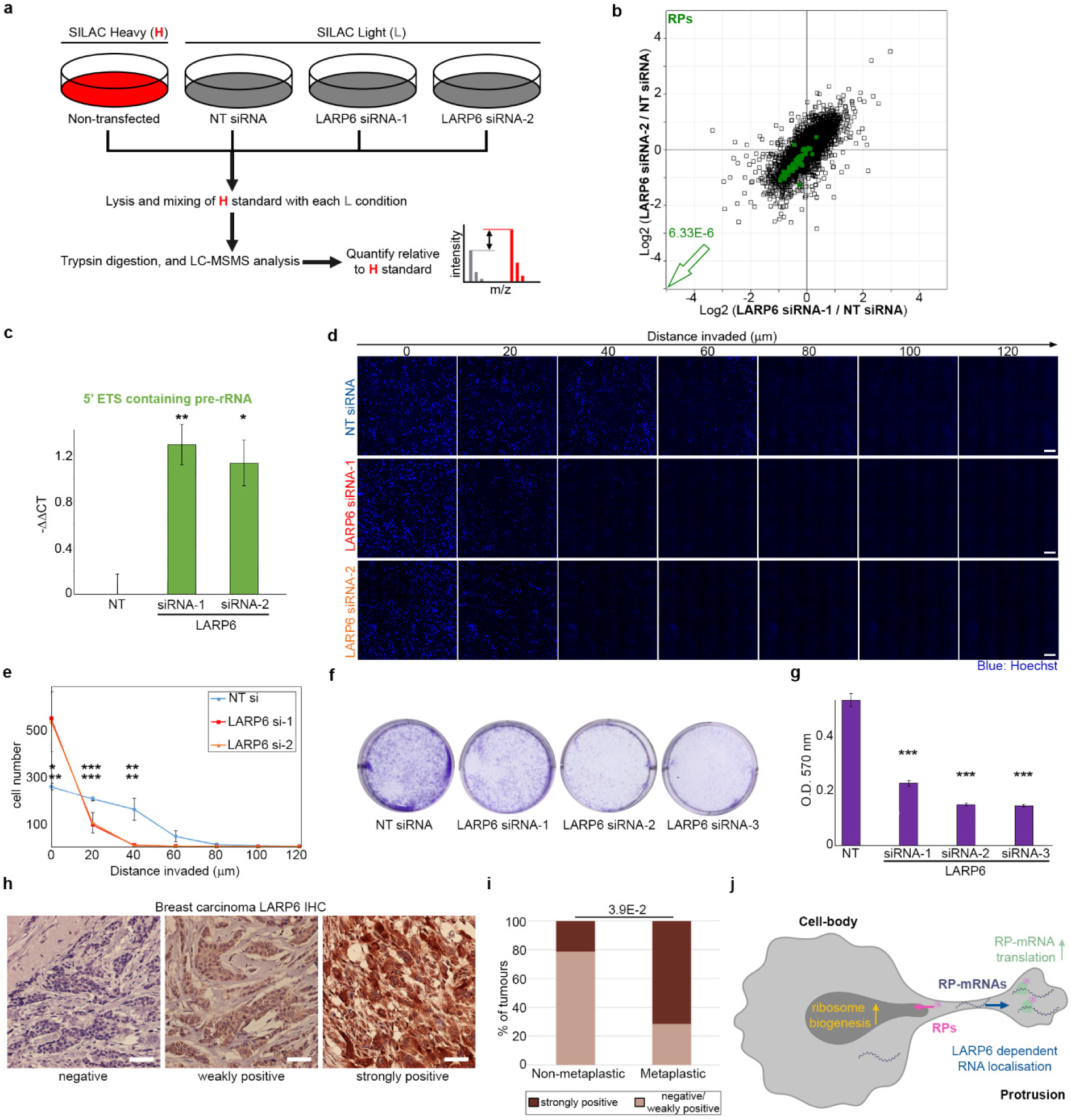
LARP6 is necessary for ribosome biogenesis, cell-proliferation, and invasion in mesenchymal-like cancer cells. **a**, Schematic representation of the quantitative proteome analysis of LARP6 depletion by SILAC. **b**, LARP6 depletion decreases RP levels (green) in actively growing MDA-MB231 cells. Changes in protein levels induced by two independent LARP6 siRNAs were quantified and plotted against each other. Benjamini-Hochberg corrected *P*-value of decrease in RP levels is reported next to the arrow. **c**, LARP6 depletion results in accumulation of pre-rRNAs. RT-qPCR analysis of 5’ETS pre-rRNA levels relative to GAPDH in MDA-MB231 cells transfected with NT control or two independent LARP6 siRNAs. \**P*<0.05; \*\**P*<0.01. **d**, LARP6 depletion hampers the ability of MDA-MB231 cells to invade through 3D Collagen-I. MDA-MB231 cells transfected as indicated for 3 days were subjected to an invasion assay through 3D Collagen-I matrix. Representative tiled confocal images of fixed, Hoechst stained cells (blue) at different migrated distances are displayed. Scale bars are 200 μm. **e**, Quantification of invaded cell numbers from (d). \**P*<0.05; \*\**P*<0.01; \*\*\**P*<0.001. **f**, LARP6 depletion hampers the proliferation of MDA-MB231 cells. MDA-MB231 cells transfected as indicated were subjected to clonogenic assays for 10 days. **g**, Quantification of experiments shown in (f). \*\*\**P*<0.001. **h**, Analysis of LARP6 expression in a panel of human breast tumors by IHC. Three distinct patterns of LARP6 expression were detected amongst the tumor samples: ‘negative’, ‘weakly positive’, and ‘strongly positive’. Scale bars are 50 μm. **i**, LARP6 strongly positive tumors are significantly enriched amongst metaplastic carcinomas. The *P*-value was calculated using Fisher’s exact test. **j**, Proposed model for LARP6 dependent regulation of ribosome biogenesis.

### Expression of LARP6 in cancer is triggered by EMT and acts to enhance protein synthesis

Since enhanced ribosome biogenesis is a common feature of most high-grade carcinomas, we wondered whether LARP6 dependent RP synthesis could be specifically upregulated in cancers to boost ribosome biogenesis. Mining a published proteomics dataset of protein expression levels in a panel of breast carcinoma cell-lines^46^ revealed LARP6 to be mainly expressed in cell-lines belonging to the mesenchymal/low Claudin subtype (Supplementary Fig. 7c). This molecular subtype is closely associated with EMT, and is primarily featured in metaplastic breast carcinomas, a rare but highly invasive form of breast cancer with poor prognosis^47^. Indeed, profiling a panel of human breast tumor tissue samples composed of both metaplastic and non-metaplastic carcinomas revealed a significant association of high LARP6 expression with metaplastic carcinomas (Fig. 5h-i). We next investigated whether the expression of LARP6 was directly triggered by EMT. Induction of EMT in epithelial-like MCF10AT cells by long-term TGFβ1 treatment, or forced expression of Snail or Twist transcription factors^47^, all resulted in a strong upregulation of LARP6 (Supplementary Fig. 7d-f). Together, these results suggest that LARP6 expression in cancer is triggered by and associated with EMT.

Due to the disproportionate upregulation of ribosome biogenesis in most high-grade cancers, there has been a great interest in developing novel strategies that can therapeutically target this pathway in clinic^2^. We next hypothesized that in cancers with strong EMT features, inhibiting LARP6 could provide a therapeutic opportunity to more specifically target ribosome biogenesis. In support of this view, induction of EMT in epithelial-like breast MCF10AT cells enhanced protein synthesis in a LARP6 dependent manner (Supplementary Fig. 7g-h). Accordingly, whilst the viability of epithelial-like MCF10AT cells was only mildly affected by LARP6 depletion, cell-viability was considerably reduced in cells that had undergone EMT (Supplementary Fig. 7i), suggesting that transformed cells which have undergone EMT are more dependent on LARP6 for supporting their protein synthesis and cell-viability.

## Discussion

Mesenchymal-like cell migration is a highly resource intensive cellular process that requires dynamic expression of large quantities of actin cytoskeletal, cell-adhesion, and extracellular matrix proteins, most of which are amongst the most abundant proteins in the proteome of mammalian cells^48^. Our results reveal a previously unknown mechanistic link between protrusion formation, which functions as the main driver of mesenchymal-like cell migration, and regulation of ribosome biogenesis, an essential cellular process that defines the protein synthetic capacity of the cells. We demonstrate that as cells protrude into their surrounding matrix, RP-mRNAs become enriched in protrusions, where they come into contact with the locally enriched translation machinery. This results in up-regulation of RP-mRNA translation. The newly synthesized RPs then travel back to the nucleus where they participate in ribosome biogenesis. Ultimately, LARP6 dependent RP-mRNA localization results in upregulation of ribosome biogenesis, leading to enhancement of overall protein synthesis (Fig. 5j). We propose that this enhancement acts as a feed-forward mechanism, enabling the cells to produce large quantities of required proteins to support invasive growth. Crucially, a recent study in mammalian gut epithelial cells also demonstrated that the subcellular localization of RP-mRNAs correlated with their translational output, although the mechanism of this localization was not defined^49^. Instead of the front-back polarity observed in mesenchymal-like cells, gut epithelial cells exhibit apical-basal polarity with distinct protein and mRNA compositions associated with each part of the polarized cell. RP-mRNAs were shown to be primarily localized to the basal portion of the cells in fasting mice, but upon feeding they translocated to the apical portion where the translation machinery was also enriched, thus leading to enhancement of their translation in an analogous feed-forward mechanism^49^. It remains to be determined whether LARP6, or another LARP family member, is similarly involved in regulation of RP-mRNAs localization in gut cells. Nevertheless, these studies collectively reveal that post-transcriptional regulation by spatial compartmentalization is a previously unappreciated mechanism in controlling RP-mRNA translation and ribosome biogenesis.

Hyperactive ribosome biogenesis is a common hallmark as well as a driver of many high-grade cancers^1, 2^. It is now evident that various anti-cancer chemotherapies function at least in part by disrupting ribosome biogenesis. Consequently, there has been a surge of interest in identifying more specific ways to target ribosome biogenesis in hope of achieving high anti-tumor activity combined with low genotoxic side effects^50^. We here show that LARP6 expression is strongly upregulated by EMT. Moreover, cells which have undergone EMT are more dependent on LARP6 for their protein synthesis and viability, collectively suggesting that LARP6 inhibition could potentially be used as a therapeutic strategy to inhibit ribosome biogenesis in carcinoma subtypes which exhibit a strong EMT signature. In addition to being more invasive, these subtypes often exhibit a greater resistance to standard chemotherapies, collectively resulting in poorer outcome^51^. Importantly, we have shown that a small molecule compound which interferes with LARP6 RNA binding activity^34^ can also inhibit RP-mRNA localization to protrusions. Although the safety, efficacy, and pharmacological properties of this specific compound may not be satisfactory for therapeutic use, our results demonstrate the plausibility of therapeutic targeting of LARP6 by small-molecule inhibitors in the context of inhibiting ribosome biogenesis in mesenchymal/EMT associated cancer subtypes.

## Experimental procedures

Material and Methods and data availability information are submitted as supplementary material.

## Author contributions

<u>F.K.M.</u> conceived the study and supervised the work. <u>M.Dermit</u> and <u>F.K.M.</u> designed the experiments, interpreted the data, and wrote the manuscript. <u>J.U.</u> and <u>S.P.B.</u> edited the manuscript. <u>M.Dodel</u> generated the MS2 reporter cell-lines, the inducible Twist and Snail cell-lines, and carried out the live cell RNA imaging, as well as the EMT vs. Parental profiling experiments. <u>F.C.Y.L.</u> and <u>M.Dermit</u> carried out the iCLIP experiments and data analysis. <u>J.U.</u> supervised all the iCLIP work. <u>M.S.A</u> performed the Quantseq 3’ FWD RNA library preparations. <u>H.S.</u> and <u>S.P.B.</u> validated the C9 compound and determined the effective dose. <u>J.L.J.</u> collected the human breast tumor tissue samples and provided the sections. All other experiments were performed by <u>M.Dermit.</u>

## Acknowledgements

We would like to thank Carme Gallego, Antonio Gentilella, Andrew Yoo, and Robert Weinberg for plasmids, and Carme Gallego, Sara Santos, and Anne Willis for advice on the experiments. All RNA-seq sequencings were carried out at the Barts and the London Genome Centre. We also wish to acknowledge the role of the Breast Cancer Now Tissue Bank in collecting and making available the human breast tissue samples used in the generation of this publication. Finally, our special thanks goes to Christopher Tape, Lovorka Stojic, and Sarah McClelland for critical reading of the manuscript. This work was funded by a Medical Research Council Career Development Award (MR/P009417/1) and a Barts Charity grant (MGU0346) to F.K.M.

