## Supplementary Information for "Subcellular mRNA localization regulates ribosome biogenesis in migrating cells"

Supplementary information for this manuscript consists of Supplementary Materials and methods, Supplementary Videos (S1 to S6), a supplementary excel file containing Datasets (S1 to S17), Supplementary Figures 1 to 7, and Supplementary references.

### Supplementary experimental procedures

#### Reagents and plasmids

The TetO-WT-L32TOP-β-Globin-12xMS2 (WT 5'TOP reporter) and TetO-MUT-L32TOP-β-Globin-12xMS2 (MUT 5'TOP reporter) constructs were a gift from Antonio Gentilella (IDIBELL, Barcelona). MCP-EGFP expression plasmid, as well as the VSV and deltaR lentiviral packaging vectors were a gift from Carme Gallego (IBMB, Barcelona). rtTA-N144 [1] was a gift from Andrew Yoo (Addgene plasmid # 66810). pTK-Twist & pTK-Snail lentiviral inducible expression plasmids [2] were a gift from Bob Weinberg (Addgene plasmids # 36977 & #36976). GFP-LARP6 expression plasmid was generated by Gateway cloning of a custom synthesized codon-optimized human LARP6 donor vector (GeneArt) into the pcDNA6.2_N-EmGFP-DEST vector (Thermo). Deletion mutation constructs of LARP6 were generated using PCR amplification of the whole LARP6 donor vector minus the targeted region, followed by recircularization via self-ligation, and gateway cloning into the pcDNA6.2_N-EmGFP-DEST vector. All LARP6 expression constructs were verified by DNA sequencing. The C9 compound was acquired as part of a compound library from ChemBridge. All other reagents used in this study are listed in Table S1.

#### Cell-lines and cell-culture

MDA-MB231, U87, and WM266.4 cells were grown in DMEM supplemented with 10% FBS, 1% Penicillin/Streptomycin. RPE cells were grown in DMEMF12, supplemented with 10% heat activated FBS, 1% Penicillin/Streptomycin; HEK293T and BJ cells were grown in DMEM supplemented with 10% heat activated FBS, 1% Penicillin/Streptomycin; PC-3M cells were grown in RPMI supplemented with 10% heat activated FBS, 1% Penicillin/Streptomycin; MCF10AT cells were grown in DMEMF12 supplemented with 5% horse FBS, 1% Penicillin/Streptomycin, 100 ng/ml cholera toxin, 20 ng/ml epidermal growth factor, 10 mg/ml insulin and 0.5 mg/ml hydrocortisone. All cells were grown in humidified incubator at 37°C with 5% $CO_2$, and routinely passaged twice per week. All cell-lines were authenticated by STR profiling (Public Health England) and were routinely checked to be mycoplasma-free by MycoAlert Plus mycoplasma detection kit.

#### 3D Collagen-I RNA-FISH

Collagen-I gel matrix was prepared as described previously [3], with slight modifications. Briefly, 5x DMEM adjusted with 0.1M NaOH and 3.7% $NaHCO_3$ was mixed with pepsinized bovine collagen-I (Advanced BioMatrix) and diluted with $dH_2O$ to 1.7 mg/mL of Collagen-I whilst on ice. The mixture was then poured into individual wells of iBidi μ-Slide with 18 wells and allowed to set at 37°C for 2h. Subsequently, the cells were plated on the top of the set matrix in complete media. After 2 days of cell invasion through the collagen-I gels, cultures were fixed with 10% Neutral Buffered Formalin (NBF) for 30 min before, further processing for RNA-FISH and antibody staining.

**3D Collagen-I Invasion assay**

3D Collagen-I invasion assays were performed as described previously [3], with some modifications. Briefly, cells were suspended in 2.3 mg/mL serum-free pepsinized bovine collagen-I (Advanced BioMatrix) to a final concentration of 100,000 cells/mL. For each condition, 200µl of the suspension was dispensed into a well of an iBidi 96-µ-plate 96 Well Blackwell plate, pre-coated with 0.2% fatty acid free BSA. 4 wells were used per condition as technical replicates. Plates were then centrifuged at 300g to collect the cells at the bottom, before incubating the plate at 37°C/10% CO2 for 2hrs to allow the Collagen to set over the cells. Subsequently, 60µl of DMEM/10% FBS was added to the top of each well to trigger invasion of the cells upward. Cells were allowed to invade overnight at 37°C/10% CO2, before being fixed and stained with addition of 8% formaldehyde in PBS, supplemented with 5µg/ml Hoechst (Thermo). The plates were then imaged on a Nikon spinning disk confocal microscope with 20X magnification, using 5x5 tile scans at 0 µm, 20 µm, 40 µm, 60 µm, 80 µm, 100 µm, and 120 µm z-planes relative to the bottom of each well.

**siRNA Transfections**

For siRNA-mediated depletions, 10,000 cells/cm2 were seeded on standard TC-treated polystyrene plates overnight. Transfections were conducted using Lipofectamine RNAiMAX (Thermo), according to manufacturer's instructions, at a final concentration of 20 nM siRNA. Cells were analyzed 72 hrs post transfection, or as indicated if otherwise. SiRNA sequences used in this study are listed in Table S1.

**Lentivirus production and transduction**

Lentiviral particles were produced in HEK293T cells by co-transfection of indicated lentiviral plus packaging VSV and deltaR vectors. 1,000,000 HEK293T cells were seeded in one well of a 6 well plate 6 hours prior to the transfection. The transfection was performed using Lipofectamine2000 (Thermo) with 2 µg of the lentivirus vector and 1 µg of each of the packaging vector, according to manufacturer's instructions. The transfection mix was then added to the medium of the cells for 12-14 hours, before removal and addition of 3 ml of fresh DMEM supplemented with 30% FCS, L-Glu, P/S, for virus production. After 24 hours, the lentivirus containing medium was harvested and passed through a 0.45um filter. Half of the supernatant was then used to reverse transduce 50,000 MDA-MB231 cells in a 6 well plate.

**MS2 reporter generation and imaging**

The MS2 reporter was generated by engineering MDA-MB231 cells to express rtTA, MCP-GFP, and the WT or MUT 5'TOP reporter constructs, via Lentiviral transduction and DNA transfection, combined with antibiotic selection, single clone selection, and FACS sorting. Briefly, rtTA-N144 lentiviral particles were produced in HEK293T cells as described above and used to reverse transduce 50,000 MDA-MB231 cells. 72 hours post transduction, the medium was exchanged with fresh DMEM containing 500 μg/ml Hygromycin B for antibiotic selection. The selection was continued whilst keeping the confluency of the cells below 50% and refreshing the selection medium every 3 days for ~2 weeks until all the cells in the negative control well were dead. Single colonies were generated using the surviving population of MDA-MB231 cells by diluting 50 cells in 10ml medium and dispersing 100 μl in each well of a 96 well plate. Verified rtTA-N144 expressing MDA-MB231 clones were then transfected with 2.5 μg of the MCP-GFP vector in a six well plate, using Lipofectamine2000 (Thermo) according to manufacturer's instructions. Two days after transfection, the cells were selected with 1500 μg/ml of G418 for ~2 week until all the cells in the negative control well were dead. 5,000,000 of the G418 selected cells were then FACS sorted to enrich for a cell population with high GFP signal, followed by generation of single colonies as mentioned above. Finally, WT or MUT 5'TOP vectors were integrated into the stable rtTA-N144 and MCP-GFP expressing MDA-MB231 clones through Lipofectamine2000 transfection as before, using 2.5 μg of the vectors. 2 days post transfection, the cells were treated with 0.5 μg/ml of Puromycin for ~10 days until all the cells in the negative control well were dead. As described before, single colonies were generated from the surviving population. A successful incorporation of the 5'TOP constructs was later verified through qPCR analysis of β-Globin expression induction following doxycycline treatment. Live-cell single molecule imaging was carried out on a Nikon spinning disk confocal microscope with 100X magnification.

**Generation of stable GFP-expressing cells**

MDA-MB-231 cells were transfected with expression constructs containing GFP-only, GFP-LARP6, or various GFP- deletion mutants of LARP6, using Lipofectamine2000 (Thermo) according to manufacturer's instructions, and selected with 10 μg/ml blasticidin for 7 days prior to FACS sorting to enrich for medium to high level GFP expressers.

**Protrusion Purification**

Cell protrusions were fractionated as described before [3], with some modifications. 10 million cells were seeded on top of 5 μg/ml collagen-I coated 75 mm polycarbonate transwell filters with 3-μm pore size (Corning), and allowed to adhere overnight without the addition of media to the bottom chamber of the transwells. The next day, the media on the top of the filter was replaced by fresh media, and protrusions were induced by addition of the same media to the bottom chamber for indicated times. For RNA-sequencing, transwells were then washed with RNase- free PBS, and RNA was purified from protrusions by shaving the bottom of the filter using a glass coverslip dipped in RLT buffer from RNeasy Mini Kit (QIAGEN). The Cell-body fraction was subsequently collected by direct addition of the RLT buffer to the top of the filter. RNA was extracted following manufacturer's instructions and quantified by Qbit RNA HS Assay Kit (Thermo). For proteomics analysis, transwells were washed by PBS, fixed with methanol for 20 min at -20°C, washed again with PBS, and the protrusions were shaved off using a glass coverslip dipped in lysis buffer (2% SDS, 100mM Tris/HCl pH 7.5). Cell-body fractions were prepared by direct addition of the lysis buffer to the top of the filter. Protein amounts were estimated by Pierce BCA Protein Assay Kit (Life Technologies) prior to sample preparation for MS.

**RNA-FISH and Immunofluorescence (IF)**

For staining of 3D invading cells, 300,000 cells grown for 2 days on 3D Collagen-I matrix filled wells of iBidi u-Slides were used. For staining of cells on 2D, 5,000 cells grown on Collagen-I coated Falcon multi-chamber slides were used. For staining of cells that protrude through transwell, 1,000,000 or 100,000 cells seeded onto 24 mm or 6.5 mm membrane inserts respectively, were used. For RNA-FISH, cells were washed with RNase-free PBS and fixed in RNase-free NBF for 30 min. The fixed cells were then washed three times with RNase-free PBS and dehydrated gradually with 50%, 70% and 100% ethanol. Cells were subsequently rehydrated gradually with 70% and 50% ethanol in RNase-free PBS, and treated with RNAScope® Protease III for 10 min prior to hybridization with pre-designed RNAScope® probes (Advanced Cell Diagnostics). All probes were then visualized using RNAscope® Fluorescent Multiplex Reagent Kit according to the manufacturer's protocol (Advanced Cell Diagnostics). If co-immunofluorescence was also conducted, samples were blocked after RNA-FISH with 10% BSA in RNase-free PBS for 20 min, and incubated with the indicated antibodies overnight at 4°C, followed by incubation with secondary antibody for 1hr at room temperature (RT). The images were acquired on Zeiss LSM 710 or 880 confocal microscopes. Imaging of protrusion and cell-body sides of transwell filters was done as described before [3], with the filters being visualized by transmitted light imaging in grey. All used antibodies in this study are listed in TABLE S1.

**RiboPuromycylation-FISH assay**

Ribopuromycylation assay was performed as described in [4], with some modifications. Briefly, transwells or slides were treated with labelling medium containing 25 µg/mL emetine plus puromycin 50 µg/ml for 5 min at 37 °C. The medium was then aspirated and slides were incubated for 20 min with ice-cold co-extraction/fixation buffer (0.015% digitonin, 5 mM MgCl2, 25 mM KCl , 0.2 M sucrose , 1x EDTA-free protease inhibitors, 1/1000 ANTI-RNase, 3% Formaldehyde, and 50 mM Tris-HCl pH 7.5, in RNase-free water). The slides were then further fixed in 10% Neutral Buffered Formalin (NBF) for 10 min at RT. The fixed cells were then washed three times with RNase-free PBS, followed by RNA-FISH and IF staining with RPL34 RNAScope® probe and anti-puromycin antibody. As control, cells were pre-treated with 2 µg/ml homoharringtonine for 15 min prior to labelling.

**Immunohistochemistry (IHC)**

A cohort of 34 Formalin-Fixed Paraffin-Embedded (FFPE) human breast carcinoma specimens consisting of 26 Invasive Ductal Carcinoma (IDC) and 7 Metaplastic breast carcinoma (MBC) samples, retrieved from the Barts Cancer Institute Breast Tissue Bank following full informed consent (ethics ref: 15/EE/0192) were analyzed by IHC. Standard 3,3'-Diaminobenzidine (DAB) method for immunostaining combined with low-pH citrate based high-pressure cooking antigen retrieval was used as reported in [5], with some modifications. Briefly, tissues were sectioned and affixed onto coated slides before being subjected to deparaffinization (two washes in xylene for 5 min) and rehydration (two washes in absolute alcohol for 2 min). Endogenous peroxidase was blocked by immersing the tissues in methanol 0.03% hydrogen peroxide in methanol twice for 5 min. Two additional washes in absolute alcohol was performed to clear out any remaining regents twice and sections were then rinsed under tap water. Subsequently, sections were heated in antigen unmasking solution (Vector labs) in a pressure cooker, reaching boiling point for 10 min and then cooled for 5 min under tap water. Sections were then dried and a hydrophobic pen was used to draw marks around tissues before transfer to wash buffer (0.2% tween in PBS). Subsequently, sections were incubated in blocking solution (2.5% bovine serum albumin and 0.2% tween in PBS) for 1 h. Next, LARP6 primary antibody (Atlas antibodies, product number: HPA049029, lot number: R58965) diluted in SignalStain® Antibody Diluent (Cell Signaling Technologies) was added and incubated overnight at 4 ° C in a wet chamber. Next day, SignalStain® Boost Detection Reagent was equilibrated to RT. Antibody solution was removed and the sections were washed with wash buffer for 3 times. Sections were then incubated with SignalStain® Boost Detection HRP rabbit reagent (Cell Signaling Technologies) in a humidified chamber for 30 min at RT, before being washed again for three times and incubation with SignalStain® DAB for 10 min, followed by immersion in water for 5 min and counterstaining with haematoxylin for 2 min. Stained sections were then dehydrated in 90% absolute alcohol for 2 min and transferred to xylene for 5 min for clearing, before mounting of a cover glass using DPX mounting medium. Specimens were then dried and visualized using an OLYMPUS BX51 microscope.

**OPP staining**

OPP staining and detection was conducted using Click-iT Plus OPP Alexa Fluor-488 Protein Synthesis Assay Kit (Thermo), according to manufacturer's instructions. Briefly, cells were treated with 10 μM OPP for 15 min at 37 °C, before being fixed with 4% formaldehyde for 15 min at RT, washed three times with PBS, and permeabilized for 5 min with 0.2% Triton X-100 in PBS. The cells were then washed three times with PBS, and the OPP labelled nascent proteins were detected using Click-iT® mediated covalent attachment of Alexa Fluor-488 azide dye. Cell were then counterstained with phalloidin (to detect cell boundaries) and NuclearMask blue (Thermo) during a 30 min incubation at RT, before three further PBS washes and imaging by confocal microscopy.

**Image analysis and statistics**

Immunofluorescence images were analyzed using ImageJ or Fiji software platforms [6]. For quantification of RNA-FISH in transwells, multi-channel color images were split, intensity levels were thresholded, followed by normalization RNA-FISH signal to the overall cell-body or protrusion areas. Protrusion and cell-body areas were defined by either CellTracker staining (Thermo), tubulin IF staining, or phalloidin labelling. Normalized protrusion to cell-body RNA-FISH values were then calculated and displayed in Log2 scale. For presentation of images, cell boundaries were marked by white dashed lines generated in the Zen blue software (Zeiss). Polarity index [7] was used as a quantification of RNA localization to the cell peripheries, and was calculated as PI= $\frac{\sqrt{(\bar{x}RNA-\bar{x}cell)2}+(\bar{y}RNA-\bar{y}cell)2}{\mathrm{Rgcell}}$, where $\bar{x}$RNA and $\bar{y}$RNA are the transcript pixel intensity positions and $\bar{x}$cell and $\bar{y}$cell are the positions for the nucleus centroid. Rgcell is the radius of gyration and it is calculated by the root-mean-square distance of all transcript pixels from the nucleus centroid of the nucleus. Co-localization analyses were performed by ComDet plugin. ComDet plugin was also used to detect and quantify the number of MCP-GFP labelled 5'TOP reporter mRNA particles from every frame image of protrusion videos. For quantification of translation, mean OPP fluorescence intensity of the cell-body images were normalized to their corresponding DAPI image intensity. LARP6 IHC staining of tumor sections were quantified using the IHC Profiler plugin [8]. This plugin allows for the color deconvolution of haematoxylin (blue) and DAB (brown) pixels. Briefly, "Nuclear Stained Image" mode was selected to find nuclei and threshold was manually set to ensure selection of malignant cells. Then, H DAB channel overlapping with malignant cells was selected for analysis on IHC Profiler using the "Cytoplasmic Stained Image" mode. IHC Profiler macro outputs of 'high positive' and 'positive' were collectively grouped as 'strongly positive', whilst the 'low positive' output was referred to as 'weakly positive'. The over-representation in metaplastic carcinomas was calculated using Fisher's exact test, with a *P*-value cut-off of 0.05. Unless stated otherwise, all *P*-values were calculated using a two-tailed homoscedastic t-test. All error bars are standard deviation, unless stated otherwise.

**Western Blotting**

Cell were lysed in 2-4% SDS, 100mM Tris/HCl pH 7.5 and sonicated with a sonicator bath (Bioruptor Pico - Rm 343) for 15 cycles. Sample concentration was adjusted with a Pierce BCA Protein Assay Kit (Thermo) before addition of NuPAGE LDS Sample Buffer (Thermo) with reducing agent and boiling at 95°C for 10 minutes. After separation on a NuPage 4%–12% Bis/Tris protein gel (Thermo), proteins were transferred to an Immobilon-P membrane (Millipore) using a standard wet transfer device. Primary antibodies were diluted in 5%BSA, PBS and incubated on the membranes at 4°C overnight followed by incubation with anti-mouse or rabbit HRP-conjugated secondary antibodies at room temperature for one hour. Membranes were then probed with Pierce ECL Plus HRP-detection reagent followed by imaging on an Amersham Imager 600. All used antibodies in this study are listed in TABLE S1.

**Colony formation and cell viability assays**

For Colony formation assay of non-targeting control or LARP6 siRNAs transfected cells, 72 hrs post transfection, 5,000 cells were seeded in 6-well TC-treated plates and allowed to grow for 10 days. Cells were then fixed with 4% formaldehyde for 30 min at 4ºC in the dark. The fixing solution was then discarded and a 0.5% crystal violet solution (0.5% w/v; 20% MeOH; 80% ddH2O) was added to the plates and incubated for 10 min at RT, before extensive washing of the plates with water. Colony images were taken with an Amersham Imager 600 machine (GE Healthcare Life Sciences). The Crystal Violet stain was then extracted with Sorenson's buffer (0.1M Na3C6H5O7; 50% EtOH; 50% ddH2O), left on agitation at 300 rpm for 30 min. Colorimetric quantification was conducted by measuring absorbance at 540 nm with a FLUOstar Omega Microplate Reader (BMG Labtech). Each biological replicate was measured in 3 technical replicates. At least 3 biological replicates were performed to calculate the average OD value. For assessment of cell viability with CellTiter-Glo® (Promega) luminescence assay, 5,000 cells/cm2 were transfected with non-targeting control or indicated LARP6 siRNAs. Three, five, or seven days post-transfection, CellTiter-Glo™ reagent was added (150ul of per well of 24 well plates). The plates were then shaken for 2 min, incubated for 10 minutes, and RLU were measured with a FLUOstar Omega Microplate Reader (BMG Labtech). Each biological replicate was measured in 3 technical replicates. At least 3 biological independent replicates were performed to calculate the average RLU value. For assessment of cell viability for C9 $IC_{50}$ calculation, WT and LARP6 KO cells were seeded 24 h prior experiment into 96 well TC-treated plates and consequently treated with C9 for 48 hrs at indicated concentrations. $IC_{50}$ measurements were calculated using MTT assay (Thermo) according to manufacturer's instructions. Readouts were normalized and $IC_{50}$ values were calculated using a non-linear regression model. Each biological replicate was performed in 4 technical replicates. At least 3 biological independent replicates were performed to calculate the average $IC_{50}$ value.

**RT-qPCR**

RT-qPCR was performed using Brilliant II SYBR® Green one-step (Agilent) with the ABI 7500 Real-Time PCR system (Applied Biosystems). The 2-ΔΔCT method was used for relative quantification of genes expression according to [9]. GAPDH was used as internal control for normalization. LARP6 expression on KDs was checked by RT-qPCR (data not shown). All primers for RT-qPCR are listed in Table S1.

**Transcriptomics Analysis**

Total RNA preps were quantified by Qubit 4 fluorimeter (Thermo). Quality of RNA was analyzed on Agilent Tapestation 4200 with High Sens. RNA ScreenTape to rule out RNA degradation (RIN ≥8). Libraries were prepared from 50-100 ng of RNA using Lexogen QuantSeq mRNA 3' end sequencing kit FW (Lexogen), according to manufacturer's instruction. Libraries were sequenced on an Illumina Nextseq 500, at Barts and the London Genome Centre. FASTQ files from QuantSeq 3' mRNA-seq data were aligned to the human reference genome using BlueBee Genomics platform. Raw read count data were uploaded into Perseus software [10] for downstream data analysis, including log2 scaling, protrusion to cell-body ratio calculation, normalization by median subtraction, Benjamini-Hochberg corrected 1D or 2D annotation enrichment analysis [11], and data visualization. Galaxy platform [12] was used to validate knockdown of isoform specific reads of LARP6 which were not differentiated by the BlueBee platform analysis.

**iCLIP**

The iCLIP method was performed as previously described in [13], with the following conditions. A total of ~ 40 million cells per biological replicate of GFP and GFP-LARP6 stably expressing MDA-MB-231 cells were irradiated once on ice with 150 mJ/cm$^2$ of UVC (254 nm), using a Hoefer Scientific UV Crosslinker. Cell pellets were lysed in iCLIP lysis buffer and diluted to a protein concentration of 1mg/ml. RNA is fragmented in lysate with RNase I at 0.4 U/ml. GFP or GFP-LARP6 was immunoprecipitated with GFP (ab290) or GFP-trap magnetic agarose beads (Chromotek). After SDS-PAGE and membrane transfer, the region corresponding to 75–200 kDa protein-RNA crosslinked complexes was excised to isolate the associated RNAs. Isolated RNAs were reverse transcribed using primers containing an experimental barcode (5nt, underlined) and UMI sequence: /5Phos/ WWW **XXXXX** NNNN AGATCGGAAGAGCGTCGTGAT /iSp18/ GGATCC /iSp18/ TACTGAACCGC. Samples were sequenced on Illumina HiSeq4000, producing 100-nt single-end reads. For data analysis, individual GFP and GFP-LARP6 iCLIP FASTQ files were uploaded onto the iMaps webserver (https://imaps.genialis.com/), which is based on the iCount package (https://icount.readthedocs.io/ en/latest/index.html), for demultiplexing and primary analysis. Reads were mapped to the GRCh38/GENCODE v27 genome. Crosslink sites were defined as the nucleotide position preceding the start of the cDNA insert (i.e. where the reverse transcription truncates). Sequencing reads arising from PCR duplication were removed by collapsing reads which map to the same crosslink site position and contain the same UMI sequence. Analysis of reproducibility of crosslink sites between biological replicates was performed by PCA, implemented in R using gene counts values. iCount group function was used to merge 6 replicates of GFP-LARP6 and 4 replicates of GFP-only individual BED files, coming from two independent biological experiments, into one BED file per condition. Reads density bar-plots were generated using the iCount summary type and subtype outputs. Metaprofile of crosslink counts normalized to total library size of the merged GFP and GFP-LARP6 replicates were plotted as RNA maps around gene start, gen end, and ORF start landmarks. Peak calling was performed using the Paraclu [14] function within iMaps, with the minimal sum of scores inside a cluster set to 10, maximal cluster size set to 200 nucleotides, and Minimal density increase set to 2. GFP-only peaks were subtracted from GFP-LARP6 peaks using bedtools intersect function in Galaxy [12] to reveal LARP6 specific binding sites. LARP6 specific target mRNAs were identified on the basis of at least having one specific LARP6 binding site. Fisher's exact test analysis of over-represented categories amongst LARP6 specific targets were performed in Perseus software [10], using an FDR cut-off of 0.02.

**Stable Isotope Labelling of Amino Acids In Cell Culture (SILAC)**

For SILAC labelling, cells were grown for at least six doublings in Lysine and Arginine free DMEM, supplemented with 10% dialyzed FBS, 1% P/S, 600mg/L Proline, in the presence of 100mg/L of either light Arginine and Lysine (for ''light'' media), medium Arginine [U-13C6] and Lysine [4,4,5,5-D4] (for ''medium'' media), or heavy Arginine [U-13C6, U-15N4] and Lysine [U-13C6, U-15N2] (for ''heavy'' media). For pulsed-SILAC, cells were grown in light SILAC media overnight, before being switched to fresh medium or heavy SILAC media for 1 to 8 hrs. After lysis, sonication, and protein concentration assessment, equal amounts of SILAC or pulsed-SILAC samples were reciprocally mixed. For pulsed-SILAC in conjugation with subcellular fractionation, cells were pulsed for 4 hrs with either heavy or medium labels, before lysis and mixing, followed by subcellular fractionation with serial solubilization.

**Mass spectrometry sample preparation and data acquisition**

Lysates, prepared in 2-4% SDS, 100mM Tris/HCl pH 7.5, were reduced with addition of 100 mM DTT and boiling at 95°C for 10 min. Filter Aided Sample Preparation (FASP) [15] was used for generation of tryptic peptides in case of SILAC/pulsed-SILAC samples. For TMT samples, isobraric Filter Aided Sample Preparation (iFASP) [16] was performed, with some modifications. Briefly, 25 μg of total protein for each sample was reduced with 50 mM Bond-Breaker TCEP Solution (Thermo) by boiling at 95°C for 10 min. Reduced samples were then diluted in UA buffer (8 M urea, 100 mM Tris HCl pH 8.5), and transferred to Vivacon 500 Hydrosart filters with a molecular cut-off of 30kDa, before being concentrated by centrifugation at 14,000 g for 20 min. Samples were then washed twice with UA buffer through cycles of buffer addition and concentration, before alkylation with addition of 10 mM iodoacetamide in UA buffer at RT for 30 min in the dark. Samples were then washed three additional times with the UA buffer, before two washes with 100 mM TEAB to reduce the urea concentration. Samples were then trypsin digested overnight at 37°C in a 600 rpm shaking thermomixer, using 40 μL of 100mM TEAB supplemented with 0.5 μg Trypsin (Sigma) per filter. Each Sample was then supplemented with 0.2 mg of a TMT label reagent at 25°C for 1 hour, followed by quenching with 5% hydroxylamine at 25°C for 30 min. Peptides were eluted by centrifugation at 14,000 g for three times, plus a further elution with 30% acetonitrile. After combining all eluates, the samples were dried with a vacuum concentrator and fractionated using Pierce™ High pH reverse-phase fractionation kit into 7 fractions, according to manufacturer's instructions. Samples were then dried with vacuum centrifugation before LC-MS/MS analysis. LC-MS/MS analysis was performed on a Q Exactive-plus Orbitrap mass spectrometer coupled with a nanoflow ultimate 3000 RSL nano HPLC platform (Thermo Fisher). Dried peptide mixtures were resuspended in 0.1% TFA, 2% Acetonitrile, and ~1-5 μg of total material was injected into the nanoflow HPLC. Samples were resolved at flow rate of 250 nL/min on an Easy-Spray 50cm X 75 μm RSLC C18 column (Thermo Fisher). Each run consisted of a 123 min gradient of 3% to 35 % of Buffer B (0.1% FA in Acetonitrile) against Buffer A (0.1% FA in LC-MS gradient water), and separated samples were infused into the MS by electrospray ionization (ESI). Spray voltage was set at 1.95 kV, and capillary temperature was set to 255°C. MS was operated in data dependent positive mode, with 1 MS scan followed by 15 MS2 scans (top 15 method). Full scan survey spectra (m/z 375-1,500) were acquired with a 70,000 resolution for MS scans and 17,500 for the MS2 scans. For TMT10plex samples, MS2 scans were acquired with 35,000 resolution. A 30 sec dynamic exclusion for fragmented peaks was enabled.

**Mass spectrometry data analysis**

MaxQuant (versions 1.5.5.1 and 1.6.3.3) was used for all mass spectrometry search and quantifications [17]. Raw data files were searched against a FASTA file of the Homo sapiens proteome, extracted from Uniprot (2016). Enzyme specificity was set to "Trypsin", allowing up to two missed cleavages. False discovery rates (FDR) were calculated using a reverse database search approach, and was set at 1%. Default MaxQuant parameters were used with some adjustments: For TMT experiments, "reporter ion MS2" type option was selected with a reporter mass tolerance of 0.01 Da. TMT 6plex or 10plex isobaric labels were selected according to the experiments. For SILAC experiments, "Match between runs" option was enabled. With the exception of pulsed-SILAC experiments, the "Re-quantify" option was also enabled. A minimum ratio count of 1 was also used for pulsed-SILAC experiments. The iBAQ calculation was also selected for nuclear, cytosol, and membrane abundance calculation of newly synthesized RPs. All downstream data analyses, such as data filtering, Log 2 transformation, ratio calculation, category annotation, 1D & 2D annotation enrichment analysis, and data visualization, were performed in Perseus software [10] (versions 1.5.5.3 and 1.6.2.3). For all annotation enrichments, GO and KEGG annotations were used, with a Benjamini-Hochberg FDR of < 0.02 applied as the cut-off in the adapted Wilcoxon Mann-Whitney test.

**Data availability**

All mass spectrometry raw files and their associated MaxQuant output files were deposited on ProteomeXchange Consortium [18] via the PRIDE partner repository (http://www.ebi.ac.uk/pride/archive/), with the following accession numbers:

1- *PXD015808* for TMT quantitative proteomics analysis of (a) protrusions and cells bodies of the normal and malignant panel of cell-lines, (b) protrusions and cells bodies of MDA-MB231 cells collected after 1 h, 2 h, 4 h, 8 h post- induction of protrusions, (c) whole lysates of non-transfected (NT) or LARP6 siRNA transfected MDA-MB231 cells grown on either closed or open pore 3 µm transwells.
2- *PXD015807* for proteomes of MDA-MB231 cells grown on either closed or open pore 3 µm transwells for either 2 or 24 hrs, as measured by TMT quantitative proteomics.
3- *PXD015795* for pulsed-SILAC quantification of translation rates between MDA-MB231 cells grown on either closed or open pore 3 µm transwells for 1, 2, 4 or 8 hrs.
4- *PXD015777* for pulsed-SILAC coupled with iBAQ quantification of protein abundances of MDA-MB231 cells in different subcellular locations, following protrusion induction.
5- *PXD015784* for SILAC of non-targeted (NT) vs LARP6 siRNA treated MDA-MB231 cells proteomics analysis.

In addition, all RNA-sequencing FASTQ files were deposited to the ArrayExpress database (http://www.ebi.ac.uk/arrayexpress) with the following accession numbers:

1- *E-MTAB-8470* for 3' mRNA-seq (FWD) sequencing of protrusion and cell-body fractions of BJ, PC-3M, RPE-1, U-87 and WM-266.4 cells.
2- *E-MTAB-8472* for 3' mRNA-seq (FWD) sequencing of protrusion and cell-body fractions of non-transfected (NT) or LARP6 siRNA transfected MDA-MB231 cells

**Table S1: List of reagents and computational tools used in this study.**

| Reagent or resource | Manufacturer | Reference |
|---|---|---|
| **RT-qPCR Primers** | | |
| h47S rRNA fw 5’ TTCGTTCGCTCGCTCGTT 3’ | SIGMA | - |
| h47S rRNA rv 5’ CAACGACACGCCCTTCTTTC 3’ | SIGMA | - |
| Hs_LARP6_1_SG QuantiTect Primer Assay | QIAGEN | QT00221445 |
| Hs_GAPDH_1_SG QuantiTect Primer Assay | QIAGEN | QT00079247 |
| **siRNAs** | | |
| SIRNA UNIV NEGATIVE CONTROL | Sigma-Aldrich | SIC001 |
| ON-TARGETplus Non-targeting Pool | Dharmacon | D-001810-10-05 |
| LARP6 siRNA-1 GGAUUCAUGGCCAUGAGA | Sigma MISSION | Hs02_00351818 |
| LARP6 siRNA-2 GCAAGAUGCUCCUGGUCUA | Sigma MISSION | Hs01_00153597 |
| LARP6 siRNA-3 CUGUGUAUAAAUACCUUCU | Sigma MISSION | Hs01_00153598 |
| LARP1 siRNA CUGACUAUGAGAUUGAUGA | Sigma MISSION | Hs01_00168468 |
| LARP1B siRNA GAGAAUGAUACACGAAGU; AGACCUGGAUCCCGGAACA; UCAAGUAAUCAACGUAAGA; GGUGGUAAUAUCCGAGGUU | Dharmacon | SMARTpool: ON-TARGETplus L-013350-02-0005 |
| LARP7 siRNA AGGAAACAGUCCGGGAUA; GUGCUAUCAAAGAGCGAAU; GCAAAGACUCAACAAGCGA; CUUAAUCAGCCUCGGGAAA | Dharmacon | SMARTpool: ON-TARGETplus L-020996-01-0005 |
| SSB siRNA GGUCGUAGAUUUAAAGGAA; GGUUAGAAGAUAAAGGUCA; GAGACCAGUAGUUUAGUAA; GGGAAGUACUAGAAGGAGA | Dharmacon | SMARTpool: ON-TARGETplus L-006877-01-0005 |

| | | |
|---|---|---|
| **Antibodies** | | |
| Rabbit polyclonal anti LARP6 | Atlas antibodies | HPA049029 |
| Mouse monoclonal anti vimentin | Abcam | ab8978 |
| Rabbit polyclonal anti GFP | Abcam | ab290 |
| GFP-Trap® | ChromoTek | gtma-20 |
| Mouse monoclonal anti E-cadherin | Cell signaling | 3195 |
| Mouse monoclonal anti TCF8/ZEB | Cell signaling | 3396 |
| Phospho-p70 S6 Kinase (Thr389) | Cell signaling | 9206 |
| 4E-BP1 | Cell signaling | 9452 |
| p70 S6 Kinase | Cell signaling | 2708 |
| α-Tubulin (DM1A) | Cell signaling | 3873 |
| α-Tubulin (11H10) | Cell signaling | 2125 |
| α-Tubulin Monoclonal Antibody | Thermo Fisher | A11126 |
| GAPDH | Novus Biologicals | NB300-21 |
| Alexa Fluor™ 488 Phalloidin | Thermo Fisher | A12379 |
| Rabbit IgG HRP linked | GE Healthcare | NA934 |
| Mouse IgG HRP linked | GE Healthcare | NA931 |
| **Chemicals** | | |
| NuPAGE LDS Sample Buffer | Life Technologies | NP0008 |
| Pierce ECL Plus Western Blotting Substrate | Life Technologies | 32132 |
| Luminata Crescendo Western HRP substrate | Fisher Scientific | 10776189 |
| Click-iT™ Plus OPP Alexa Fluor™ 488 Protein Synthesis Assay | Thermo Fisher | C10456 |
| Crystal violet | Sigma-Aldrich | C6158 |
| DTT | VWR | M109 |
| Iodoacetamide | VWR | 786-228 |
| Blasticidin S HCl | Life Technologies | R21001 |
| TGF-β1 human | SIGMA | H8541 |
| Emetine | SIGMA | E2375 |
| Harringtonine | SIGMA | SML1091 |
| Puromycin | SIGMA | P9620 |
| ANTI-RNase (15-30 U/µL) | Life Technologies Ltd Invitrogen Division | AM2692 |
| RNase A, DNase and protease-free (10 mg/mL) | Life Technologies Ltd | EN0531 |
| AZD8055 | Selleckchem | S1555 |
| Torin | Selleckchem | |
| Everolimus | Selleckchem | S1120 |
| HCS NuclearMask™ Blue Stain | ThermoFisher | H10325 |
| SignalStain® Antibody Diluent | Cell signaling | 8112S |
| SignalStain® DAB Substrate Kit | Cell signaling | 8059P |
| SignalStain® Boost IHC Detection Reagent (HRP, Rabbit) | Cell signaling | 8114P |

| | | |
|---|---|---|
| Antigen Unmasking Solution, Citric Acid Based | Vector Laboratories | H-3300 |
| DPX new | Merck | 100579 |
| RNAscope® Fluorescent Multiplex Reagent Kit (320850) | Advanced Cell Diagnostics Srl | 320850 |
| Hs-RPL34 targeting 24-702 of NM_000995.4 | Advanced Cell Diagnostics Srl | 504031 |
| Hs-RPS7 targeting 2-521 of NM_001011.3 | Advanced Cell Diagnostics Srl | 504211 |
| Hs-RPLP2 | Advanced Cell Diagnostics Srl | 511391 |
| Hs-RPL22 | Advanced Cell Diagnostics Srl | 435271 |
| Hs-RPS21 | Advanced Cell Diagnostics Srl | 511381 |
| Hs-ITGB4 | Advanced Cell Diagnostics Srl | 300031 |
| **TC products** | | |
| 75 mm Transwell with 3.0 µm pore polycarbonate membrane insert | Corning | 3420 |
| 24 mm Transwell with 3.0 µm pore polycarbonate membrane insert | Corning | 3414 |
| 6.5 mm Transwell with 3.0 µm pore polycarbonate membrane insert | Corning | 3415 |
| iBiDi u-Slide 18 Well flat, ibiTreat, Tissue Culture Treated, Sterile | Thistle Scientific | 81826 |
| CellTracker™ Green CMFDA Dye | ThermoFisher | C7025 |
| CellTracker™ Orange CMTMR Dye | ThermoFisher | C2927 |
| Falcon™ Chambered Cell Culture Slides | Thermo Fisher | 354114 |
| Collagen I | Advanced BioMatrix | 5005 |
| Lipofectamine 2000 Transfection Reagent-1.5 mL | Life Technologies | 11668019 |
| Lipofectamine™ RNAiMAX Transfection Reagent | Thermo Fisher | 13778150 |
| Opti-MEM I Reduced Serum Medium-100 mL | Thermo Fisher | 31985062 |
| Gibco™ DMEM w/High Glucose and w/o Glutamine, Lysine and Arginine | Fisher | 12817552 |
| **Commercial reagent sets** | | |
| TMTsixplex™ Isobaric Label Reagent Set | Thermo Fisher | 90061 |
| TMT10plex™ Isobaric Label Reagent Set | Thermo Fisher | 90110 |
| Subcellular Protein Fractionation Kit | Thermo Fisher | 78840 |
| Pierce High pH Reversed-Phase Peptide Fractionation Kit | Life Technologies | 84868 |
| QuantSeq mRNA 3' end sequencing kit | Lexogen | SKU: 015.24 |
| CellTiter-Glo® Luminescent Cell Viability Assay | Promega | G7571 |
| MTT Cell Viability Assay | Thermo Fisher | M6494 |
| High Sens. RNA ScreenTape Sample Buffer | Agilent | 5067-5580 |
| High Sensitivity RNA ScreenTape | Agilent | 5067-5579 |
| **Software and Algorithms** | | |
| Maxquant | N/A | https://www.biochem.mpg.de/5111795/maxquant |
| Perseus | N/A | https://www.biochem.mpg.de/5111810/perseus |
| BlueBee | N/A | https://www.bluebee.com/ |
| Galaxy | N/A | https://usegalaxy.org/ |

| | | |
|---|---|---|
| GradPad PRISM | N/A | https://www.graphpad.com/scientific-software/prism/ |
| iMAPS | N/A | https://imaps.genialis.com/iclip |

### Supplementary Videos

**Video S1:** Time-lapse analysis of WT 5'TOP MS2 reporter MDA-MB231 cells without doxycycline. 200ms frame images were taken for 10 seconds at 100X magnification, showing diffuse localization of MCP-GFP.

**Video S2:** Time-lapse analysis of WT 5'TOP MS2 reporter MDA-MB231 cells with doxycycline (2μg/mL for 12 hrs). 200ms frame images were taken for 10 seconds at 100X magnification, showing MCP-GFP bound cytoplasmic WT 5'TOP mRNA particles.

**Video S3:** Time-lapse analysis of MUT 5'TOP MS2 reporter MDA-MB231 cells without doxycycline. 200ms frame images were taken for 10 seconds at 100X magnification, showing diffuse localization of MCP-GFP.

**Video S4:** Time-lapse analysis of MUT 5'TOP MS2 reporter MDA-MB231 cells with doxycycline (2μg/mL for 12 hrs). 200ms frame images were taken for 10 seconds at 100X magnification, showing MCP-GFP bound cytoplasmic MUT 5'TOP mRNA particles.

**Video S5:** Time-lapse analysis of protrusions of WT 5'TOP MS2 reporter MDA-MB231 cells with doxycycline (2μg/mL for 12 hrs). Cells were grown on transwell filters and induced to form protrusions for 2 hrs prior to imaging. 200ms frame images were taken for 3 seconds at 100X magnification from the bottom of transwell filters, showing MCP-GFP bound WT 5'TOP mRNA particles within protrusions.

**Video S6:** Time-lapse analysis of protrusions of MUT 5'TOP MS2 reporter MDA-MB231 cells with doxycycline (2μg/mL for 12 hrs). Cells were grown on transwell filters and induced to form protrusions for 2 hrs prior to imaging. 200ms frame images were taken for 3 seconds at 100X magnification from the bottom of transwell filters, showing no MCP-GFP bound MUT 5'TOP mRNA particles within protrusions.

### Supplementary Figures

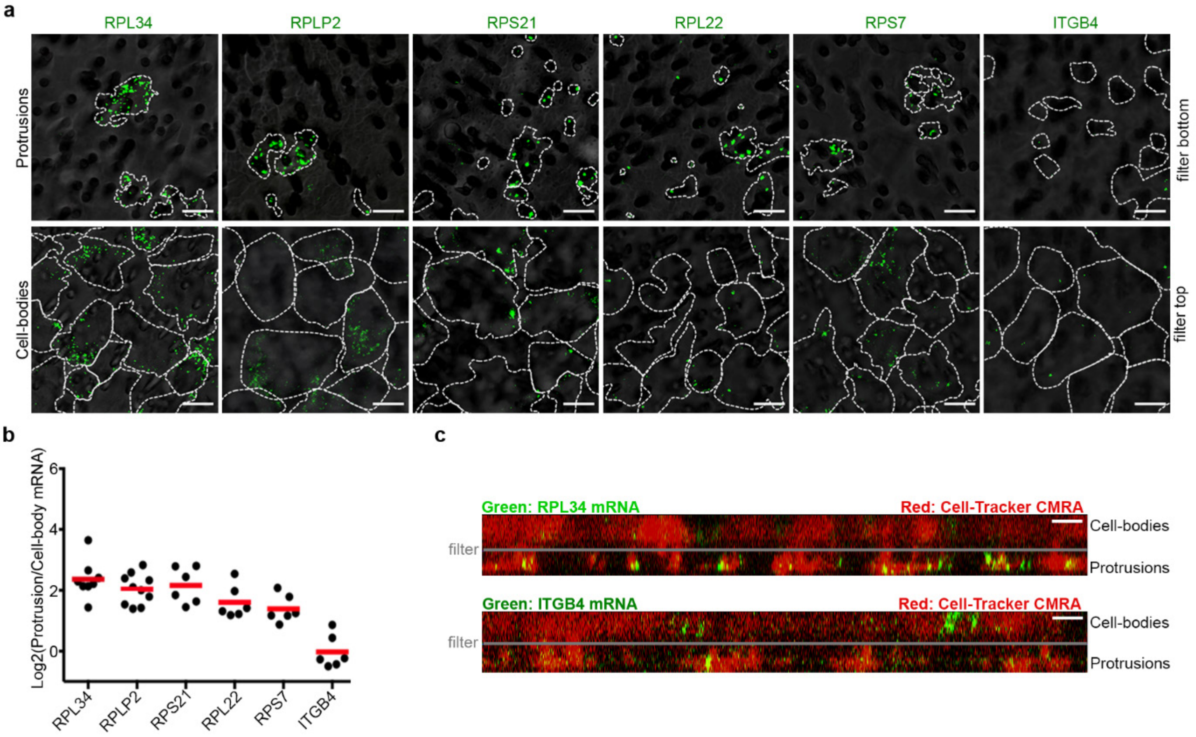

**Supplementary Fig. 1: RP-mRNAs localize to protrusions. a,** Validation of RP-mRNA localization to protrusions of MDA-MB231 cells by RNA-FISH. Representative confocal images of protrusions and cell-body filter sides stained with RNA-FISH probes against the indicated mRNA (Green). Cell boundaries (dash-lines) were defined by co-staining of the cells with anti-tubulin antibody or CellTracker. The filters (grey) were visualized by transmitted light microscopy. **b,** Quantification of protrusion to cell-body RNA-FISH ratio values from experiments shown in (a). Each data-point represents an independent large field of view image. **c,** Protrusions formed through transwell filters are enriched in RPL34 mRNA whereas cell-bodies are enriched with ITGB4 mRNA. Cross-section views of confocal Z-stack images of MDA-MB231 cells protruding through 3-µm transwell filters. Cells were stained with CellTracker (red), fixed and analyzed with FISH for RPL34 mRNA and ITG4B mRNA (green). Grey lines mark the position of the polycarbonate filter.

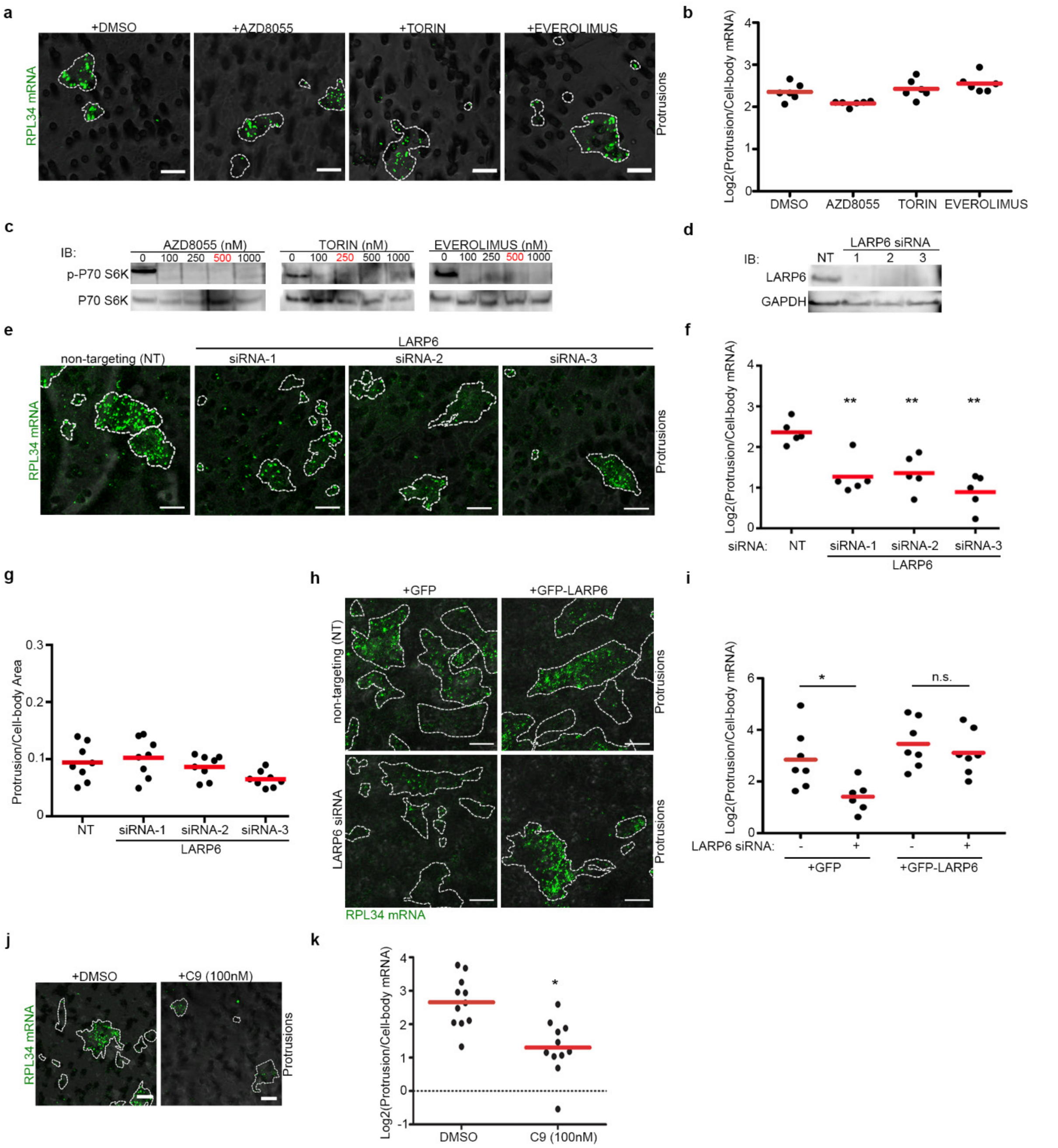

**Supplementary Fig. 2: LARP6 localizes RP-mRNAs to protrusions. a,** Localization of RP-mRNAs to protrusions is independent of mTORC1. Representative RNA-FISH images of RPL34 mRNA (green) in protrusions of mock treated vs. mTORC1 inhibitor treated (2 hrs) MDA-MB321 cells. Protrusion boundaries (dash-lines) were defined by co-staining of with anti-tubulin antibody. **b,** Quantification of protrusion to cell-body ratio values for RPL34 mRNA from experiments shown in (a). Each data-point represents a large field of view image. **c,** Validation of mTORC1 inhibitors by immunoblotting (IB). MDA-MB231 cells were treated for 2 hrs with indicated doses of mTORC1 inhibitors and assessed by IB for the phosphorylation status of P70 S6 Kinase. Inhibitor concentrations highlighted in red were used for the localization analyses in (a). p-P70 S6K: Phospho-p70 S6 Kinase (Thr389). **d,** Validation of siRNA mediated LARP6 KD by immunoblotting (IB). MDA-MB231 cells were transfected with the indicated siRNAs for 72 h before IB with the indicated antibodies. **e,** Validation of LARP6 effect on RP-mRNA localization by 3 independent siRNAs. Representative images of RPL34 mRNA in protrusions of MDA-MB231 cells (green) transfected with NT control or independent LARP6 siRNAs. Protrusion boundaries (dash-lines) were defined by co-staining with anti-tubulin antibody. The transwell filters (grey) were visualized by transmitted light microscopy. **f,** Quantification of RPL34 mRNA enrichment in protrusions from experiments shown in (e). Each data-point represents a large field of view image. **P<0.01. **g,** Protrusion formation is not affected by LARP6 depletion. Relative protrusion/cell-body areas from experiments shown in (e). **h,** Expression of an siRNA resistant LARP6 construct rescues RP-mRNA localization to protrusions. Representative images of RPL34 mRNA (green) in protrusions of MDA-MB231 cells stably expressing GFP or GFP-LARP6, transfected with NT control or an siRNA against 3'UTR of endogenous LARP6. Protrusion boundaries (dash-lines) were defined by co-staining with anti-tubulin antibody. The transwell filters (grey) were visualized by transmitted light microscopy. **i,** Quantification of protrusion to cell-body ratio values for RPL34 mRNA from experiments shown in (h). Each data-point represents a large field of view image n.s.: non-significant; *P<0.05. All scale bars are 10 μm. **j,** Treatment of transwell protruding cells with a specific LARP6 inhibitor named C9 blocks RP-mRNA localization to protrusions. Representative RNA-FISH images of RPL34 mRNA in protrusions of MDA-MB231 that were allowed to protrude through transwells for 2 hrs in presence of DMSO or C9. Protrusion boundaries (dash-lines) were defined by co-staining with anti-tubulin antibody. Scale bars are 10 μm. **k,** Quantification of RPL34 mRNA enrichment in protrusions from experiments shown in (j). Each data-point represents a large field of view image. *P<0.05.

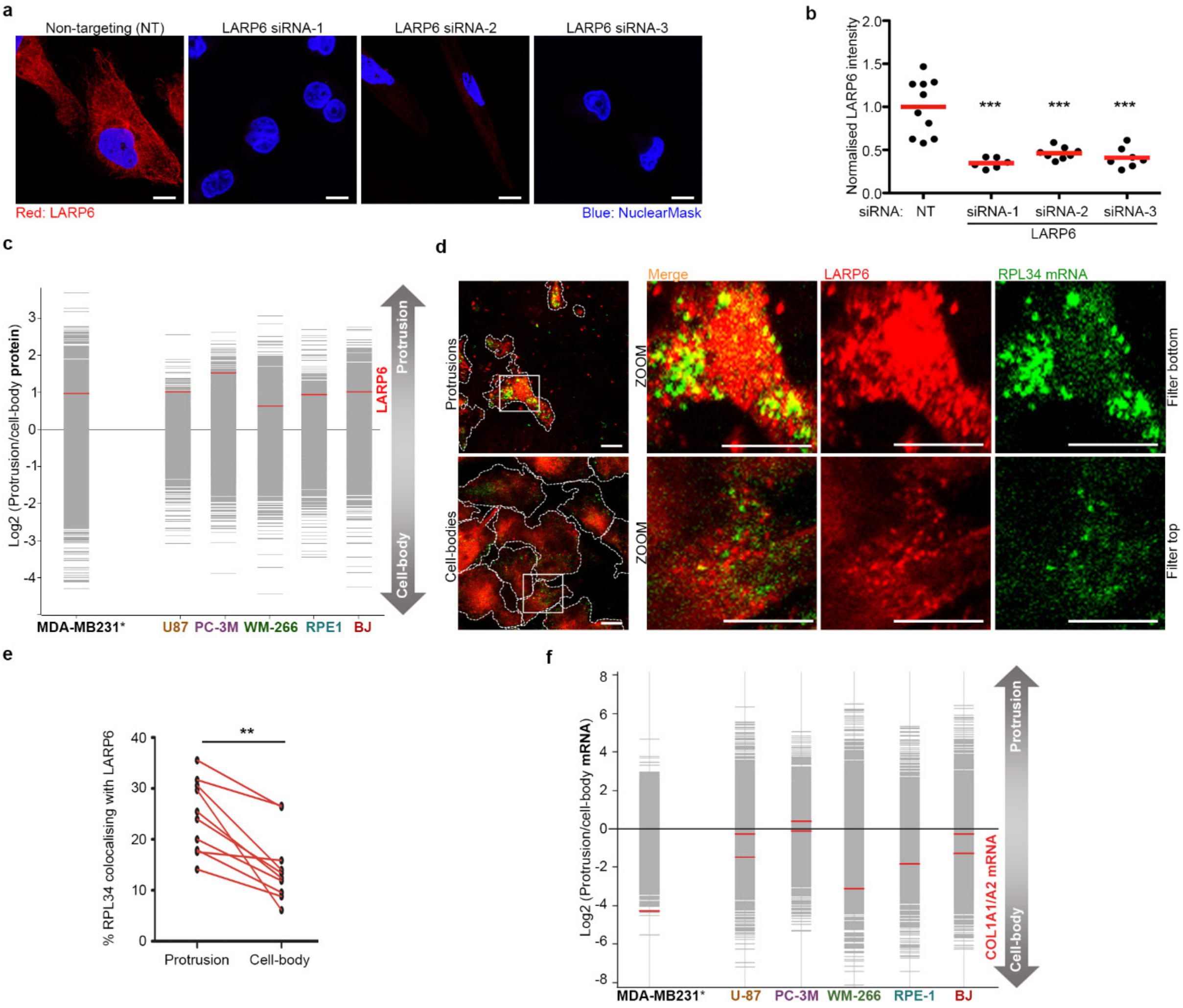

**Supplementary Fig. 3: LARP6 co-localizes with RP-mRNAs in protrusions. a,** Validation of LARP6 antibody specificity by immunofluorescence. Representative immunofluorescence images of LARP6 IF staining in MDA-MB231 cells transfected with indicated siRNAs for 72 h. **b,** Quantification of LARP6 fluorescence intensity relative to the mean NT control value from experiments shown in (a). Each data-point represents a large field of view image. ***$P$<0.01. **c,** LARP6 protein is enriched in protrusions of all the cell-lines investigated. Protrusion to cell-body protein levels from all profiled cell-lines, measured by TMT proteomics, were plotted with LARP6 being highlighted in red. *MDA-MB231 data was obtained from[3]. **d**, LARP6 co-localizes with RP-mRNAs in protrusions. Representative RNA-FISH and IF co-staining images of RPL34 mRNA (green) and LARP6 (red) in protrusions and cell-bodies of MDA-MB231 cells. Cell boundaries (dash-lines) were defined from co-staining with anti-tubulin antibody. The white rectangle marks the zoom area. **e,** Quantification of the % of co-localization of RPL34 mRNA with LARP6 granules in corresponding protrusion and cell-body images from experiments shown in (d). Red lines connect values of protrusion and body images from same field of view confocal scans. **P<0.01. **f,** Collagen-I mRNAs are mainly enriched in the cell-bodies. Protrusion to cell-body mRNA distributions from all profiled cell-lines were plotted with COL1A1/A2 mRNAs being highlighted in red. *MDA-MB231 data is obtained from[3].

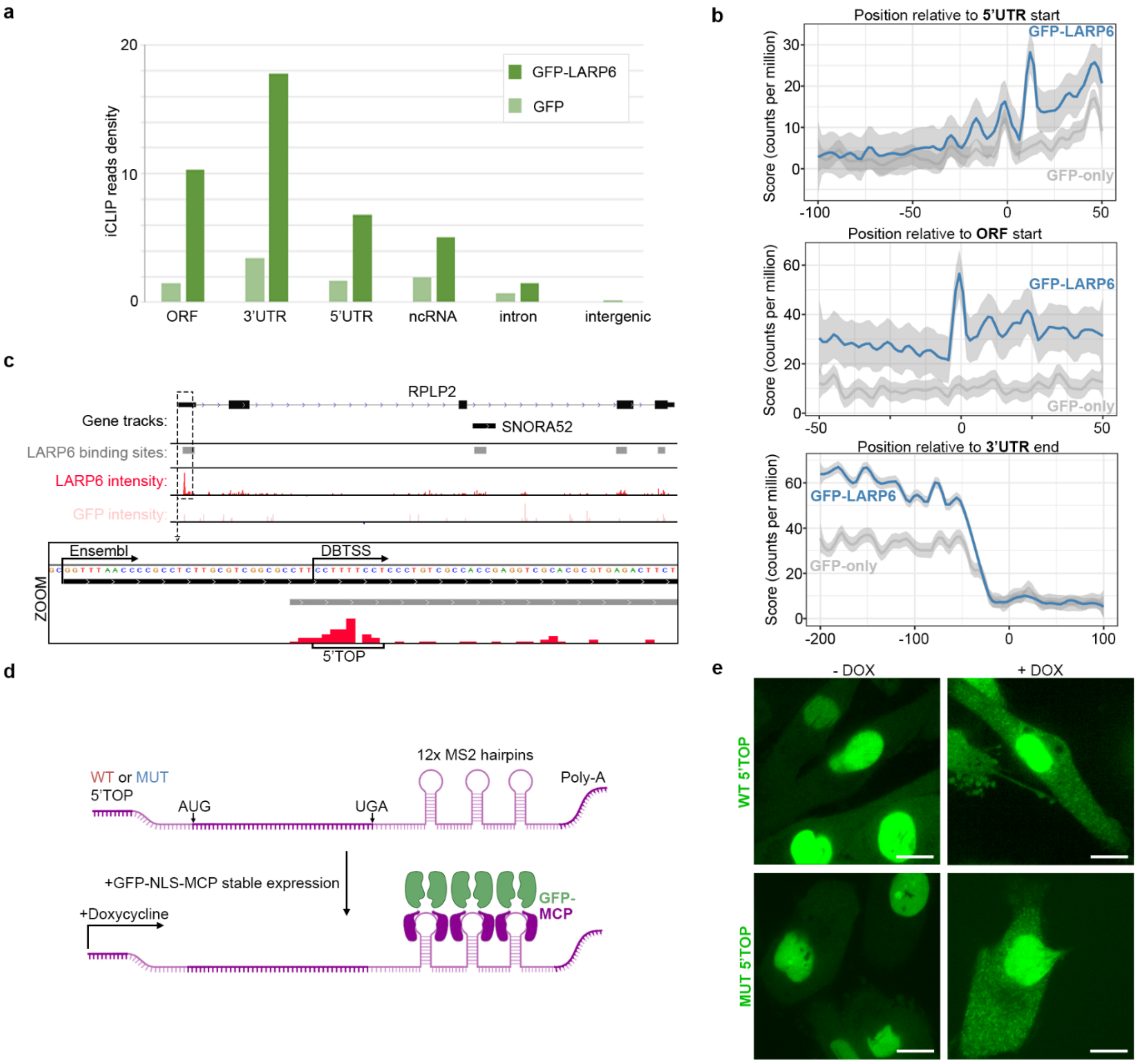

**Supplementary Fig. 4: LARP6 directly binds RP-mRNAs. a,** iCLIP reads densities corresponding to different genomic regions from GFP-only vs. GFP-LARP6 iCLIP experiments. **b,** Metaprofile plots of GFP vs. GFP-LARP6 iCLIP crosslink sites at the indicated aligned genomic regions showing LARP6-specific bindings to 5'UTR, ORF, and 3'UTR. **c,** An example genomic view of LARP6 specific binding sites after peak calling (grey) in an RP-mRNA (RPLP2), along with read intensities for GFP-LARP6 and GFP only iCLIP runs. Four distinct LARP6 binding sites are mapped to the RPLP2 locus: two mapping to the ORF region, one to RPLP2 3rd intron which is also annotated as small nucleolar RNA SNORA52, and one to the 5'UTR. Inset: zoomed view of RPLP2 5'UTR showing the LARP6 binding site overlapping with the 5'TOP. Note that for most RP-mRNAs, the Ensembl annotation of TSS is further upstream of the more accurately annotated TSS via TSS-seq in DBTSS[19]. **d,** Schematic representation of the MS2 reporter system for live cell monitoring of 5'TOP mediated RNA localization. MDA-MB231 cells were engineered to stably co-express a GFP tagged MS2 Coat Protein (GFP-MCP) with doxycycline inducible WT or MUT 5'TOP mRNA reporter constructs containing the ORF region of β-globin followed by twelve MS2 hairpin repeats in the 3'UTR[20]. Upon doxycycline induction, the reporter mRNA molecules are expressed, with each molecule binding to multiple GFP-MCP proteins that allows their detection by live cell fluorescence microscopy. **e,** Validation of the MS2 reporter system described in (e). GFP-MCP exhibits a diffuse cytosolic staining with enrichment in the nucleus (due to presence of an NLS signal) in the absence of doxycycline (-DOX). Cytoplasmic RNA particles can be observed in the presence of doxycycline (+DOX), each corresponding to a single molecule of WT or MUT 5'TOP MS2 reporter mRNA. All scale bars are 10 µm.

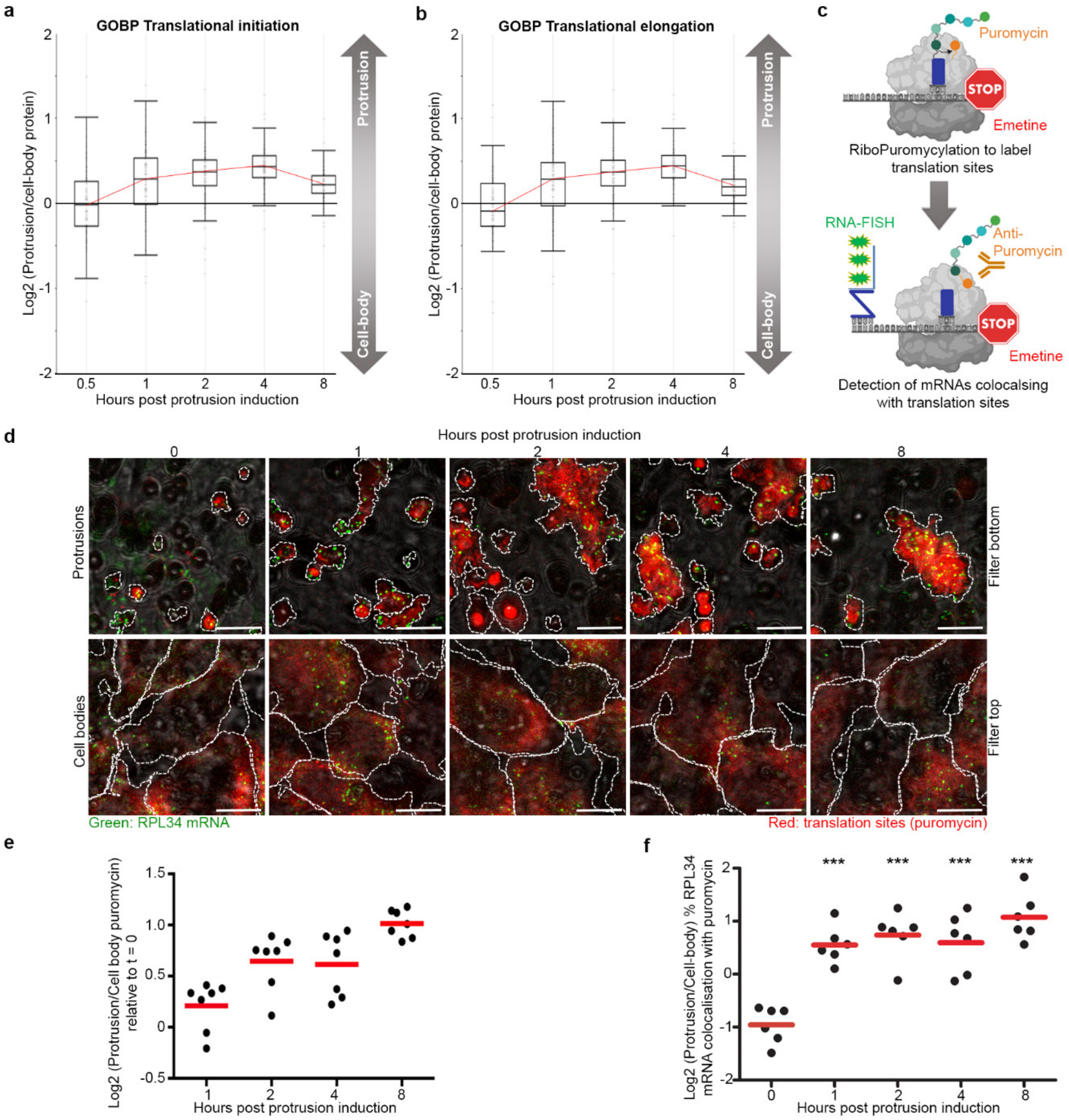

**Supplementary Fig. 5: Localization of RP-mRNAs to protrusions enhances their translation. a,** Time-course changes in the distribution of proteins annotated in GOBP as 'translational initiation' between protrusions and cell-bodies of MDA-MB231 cells following protrusion induction. **b,** Time-course changes in the distribution of proteins annotated in GOBP as 'translational elongation' between protrusions and cell-bodies of MDA-MB231 cells following protrusion induction. **c,** Schematic representation of the Ribopuro-FISH assay. Ribopuromycilation (RPM)[4] coupled with RNA-FISH can reveal active sites of translation and quantify how much an mRNA of interest is associated with such sites. **d,** RP-mRNAs are associated with active sites of translation in protrusions. Representative Ribopuro-FISH images of RPL34 mRNA (green) and puromycin (red) in protrusions and cell-bodies of MDA-MB231 cells at the indicated time points post protrusion induction. Cell boundaries (dash-lines) were defined by co-staining with anti-tubulin antibody. All scale bars are 10 μm. **e,** Translation sites in protrusions relative to the cell-bodies increases over time. Quantification of puromycin staining intensities in protrusions relative to cell-bodies, from experiments shown in (d). Each data-point represents a large field of view image normalized to time zero. **f,** Association of RPL34 mRNAs with active sites of translation is higher in protrusions than cell-bodies. Quantification of % RPL34 mRNA co-localization with puromycin in protrusions relative to cell-bodies from experiments shown in (d). Each data-point represents a field of view image. ***$P<0.001$.

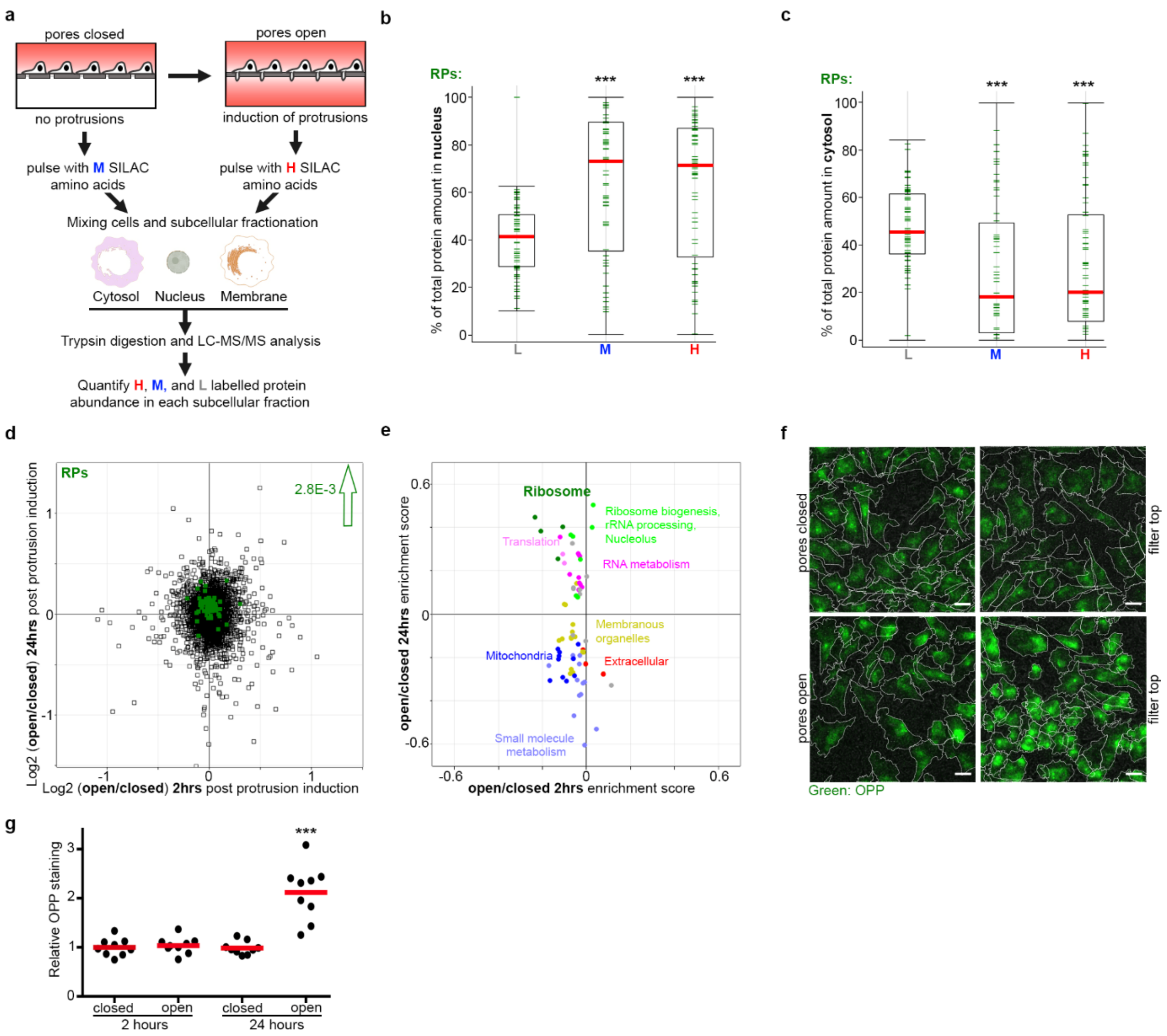

**Supplementary Fig. 6: Protrusion induction enhances RP-mRNA translation and ribosome biogenesis. a,** Schematic representation of the experimental outline for pulsed-SILAC mediated assessment of subcellular distributions of nascent proteins following protrusion induction. Absolute abundances of old unlabeled (L), or newly synthesized medium (M) and heavy (H) SILAC labelled proteins in each subcellular compartment were measured by iBAQ, in presence or absence of protrusions, and used to calculate the % of labelled protein in each compartment. **b,** Newly synthesized RPs accumulate in the nucleus. Box plot of the % of old and new RPs in the nuclear fraction of MDA-MB231 cells. Old RPs (L), nascent RPs synthesized under basal condition without protrusions (M), and nascent RPs synthesized under protrusion induced condition (H) were distinguished by their SILAC labelling state. Error bars are min-max range. ***$P$<0.001. **c,** Newly synthesized RPs are depleted from cytosol. Box plot of the % of old and new RPs in the cytosolic fraction of MDA-MB231 cells. % of old and nascent RPs in the cytosol were quantified and plotted as in (b). ***$P$<0.001. **d,** Protrusion formation enhances total RP levels. Changes in protein levels between cells with or without protrusions following 2 or 24 hrs of protrusion formation were quantified by TMT proteomics. RP levels (green) increase upon protrusion formation for 24 but not 2 hrs. Benjamini-Hochberg corrected $P$-value of the shift in RP levels is reported next to the arrow. **e,** 2D-annotation enrichment analysis of data shown in (d). Each data point represents a GO or KEGG protein category, with similar categories being highlighted by similar colors. Categories comprised of RPs (dark green), as well as ribosome biogenesis/rRNA processing-related (light green) and translation-related proteins (light pink) are upregulated upon 24 hrs of protrusion formation. **f,** Protrusion formation enhances overall protein synthesis. Transwell seeded MDA-MB231 cells were either prevented from protruding through pores (pores closed), or allowed to form protrusions (pores open), for 2 or 24 hrs, before labelling with OPP for 15 mins. Representative images of the cells from top of the filters are displayed. Scale bars are 20 µm. **g,** Quantification of normalized OPP levels from experiments shown in (f). Each data-point represents a large field of view image. ***P<0.001.

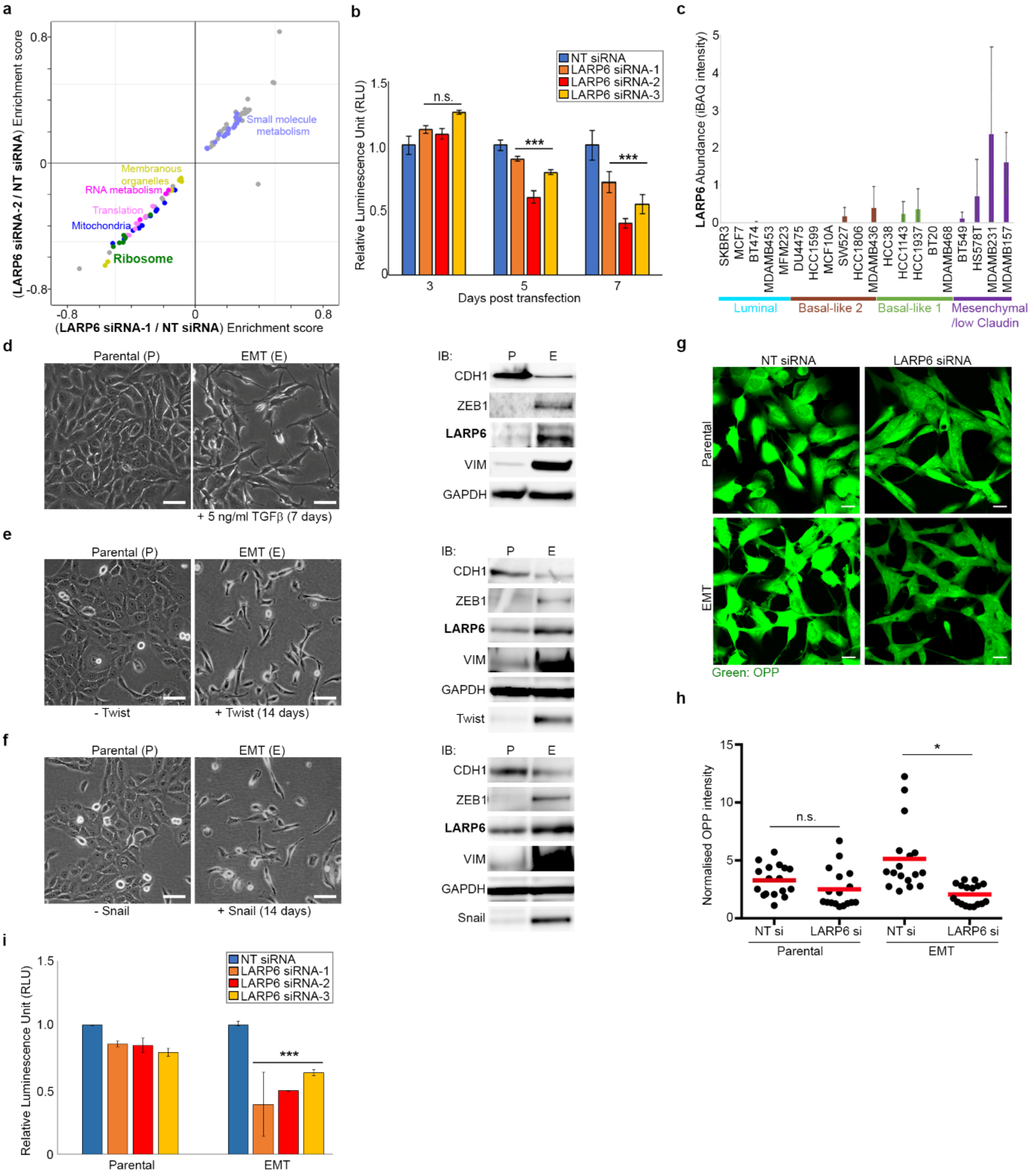

**Supplementary Fig. 7: LARP6 expression is triggered by EMT and enhances ribosome biogenesis and cell-proliferation. a,** 2D-annotation enrichment analysis of data shown in Fig 5b. Each data point represents a protein category inferred from GO and KEGG, and similar categories are highlighted by similar colors. Categories of proteins comprised of RPs (green), translation-related (light pink), and RNA metabolism-related (pink) are all significantly downregulated upon LARP6 depletion by two independent siRNAs. **b,** Cell viability of MDA-MB231 cells quantified by CellTiter-Glo assay at indicated time-points after transfection with indicated siRNAs. Averages were calculated from 3 biological replicates, each performed in 3 technical replicates. ***$P$<0.001. **c,** LARP6 protein is mostly detectable amongst cell-lines belonging to the mesenchymal/low Claudin molecular subtype of breast cancer. LARP6 absolute protein abundances, calculated by iBAQ, were extracted from[21] and plotted across the profiled cell-lines belonging to the different molecular subtypes of breast cancer. **d,** Induction of EMT by human TGFβ1 upregulates LARP6. Left: morphology of MCF10AT cells, following mock treatment or TGFβ1 (5ng/ml) treatment for 7 days. Scale bars are 50 µm. Right: IB analysis of EMT markers (CDH1, ZEB1, VIM) and LARP6, on the cells shown on the left. GAPDH was used as loading control. P: Parental; E: EMT. **e,** Induction of

EMT by overexpression of Twist upregulates LARP6. Left: morphology of MCF10AT cells stably harboring a doxycycline inducible Twist construct, with or without doxycycline treatment (1μg/ml) for 14 days. Scale bars are 50 μm. Right: IB analysis of EMT markers (CDH1, ZEB1, VIM), Twist, and LARP6, on the cells shown on the left. GAPDH was used as loading control. P: Parental; E: EMT. **f,** Induction of EMT by overexpression of Snail upregulates LARP6. Left: morphology of MCF10AT cells stably harboring a doxycycline inducible Snail construct, with or without doxycycline treatment (1μg/ml) for 14 days. Scale bars are 50 μm. Right: IB analysis of EMT markers (CDH1, ZEB1, VIM), Snail, and LARP6, on the cells shown on the left. GAPDH was used as loading control. P: Parental; E: EMT. **g,** EMT enhances overall protein synthesis in a LARP6 dependent manner. MCF10AT parental and EMT pairs from (d) were treated with NT control or LARP6 siRNAs for 72 hrs before being subjected to OPP staining. **h,** Quantification of OPP staining from experiments shown in (g). Each data-point corresponds to a large field of view image. n.s.: non-significant; **P*<0.05. **i,** Cells that have undergone EMT are more sensitive towards loss of LARP6. Cell-viability of MCF10AT Parental and EMT pairs from (d), following transfection with indicated siRNAs for 72 hrs, was quantified by CellTiter-Glo assay. Averages were calculated from 3 biological replicates, each measured in 3 technical replicates. ****P*<0.001.
